# A revised mechanism of action of hyperaldosteronism-linked mutations in cytosolic domains of GIRK4 (KCNJ5)

**DOI:** 10.1101/866202

**Authors:** Boris Shalomov, Reem Handklo-Jamal, Haritha P. Reddy, Neta Theodor, Amal K. Bera, Nathan Dascal

## Abstract

G-protein gated, inwardly rectifying potassium channels (GIRK) mediate inhibitory transmission in brain, heart, and adrenal cortex. GIRK4 (*KCNJ5*) subunits are abundant in the heart and adrenal cortex. Multiple mutations of *KCNJ5* cause primary aldosteronism (PA). According to a leading concept, mutations in the pore region of GIRK4 cause loss of K^+^ selectivity; the ensuing Na^+^ influx depolarizes zona glomerulosa cells and activates voltage gated Ca^2+^ channels, inducing hypersecretion of aldosterone. The concept of selectivity loss has been extended to mutations in cytosolic domains of GIRK4 channels, remote from the pore region. We expressed GIRK4_R52H_, GIRK4_E246K_, and GIRK4_G247R_ mutants in *Xenopus* oocytes and human adrenocortical carcinoma cell line (HAC15). Whole-cell currents of heterotetrameric GIRK1/4_R52H_ and GIRK1/4_E246K_ (but not GIRK1/4_G247R_) channels were greatly reduced compared to GIRK1/4_WT_. Nevertheless, all heterotetrameric mutants retained full K^+^ selectivity and inward rectification. When expressed as homotetramers, only GIRK4_WT_, but none of the mutants, produced whole-cell currents. Confocal imaging, single channel and Förster Resonance Energy Transfer (FRET) analyses showed: 1) reduction of membrane abundance of all mutated channels, especially as homotetramers, 2) impaired interaction with Gβγ subunits, and 3) reduced open probability of GIRK1/4_R52H_. VU0529331, a GIRK4 opener, activated homotetrameric GIRK4_G247R_ channels, but not GIRK4_R52H_ and GIRK4_E246K_. Our results suggest impaired gating (GIRK4_R52H_) and expression in plasma membrane (all mutants). We suggest that, contrary to the previously proposed mechanism, R52H and E246K mutants are loss-of-function rather than gain-of-function/selectivity-loss mutants. Hence, GIRK4 openers may be a potential course of treatment for patients with cytosolic N- and C-terminal mutations.

**Significance Statement:** Mutations in KCNJ5 gene, which encodes for the GIRK4 subunit of G-protein inwardly rectifying K+ channels, are the main cause of primary aldosteronism, a major contributor to secondary hypertension. We report that three mutations in the cytosolic domain of GIRK4 cause loss-of-function, contrary to the prevailing concept that these mutations cause loss of selectivity and subsequent depolarization, i.e. essentially gain-of-function. Our findings correct the existing misconception regarding the biophysical mechanism that impairs the channel function, and may provide indications for future personalized treatment of the disease.

## Main Text

### Introduction

G-protein gated, inwardly rectifying potassium channels (GIRK; Kir3) mediate inhibitory signaling via G-protein coupled receptors (GPCR) in the brain, heart, and adrenal cortex. There are four GIRK subunits (GIRK1-4) forming homo-(GIRK2, GIRK4) or heterotetrameric channels (GIRK1/2, GIRK1/3, GIRK1/4, GIRK2/3). Gβγ dimers are the main gating factor of GIRK channels (1-3); upon GPCR activation, Gβγ dissociates from Gα_i/o_, binds to the channel and activates it (4). Like in all inward rectifiers, in GIRKs the outward current is smaller than the inward, due to the occlusion of the cytoplasmic pore by Mg^2+^ and polyamines (5). GIRK4 subunits, encoded by *KCNJ5* gene, are mostly abundant in heart and adrenal cortex, forming GIRK4 homo-and/or GIRK1/4 (*KCNJ3/KCNJ5*) heterotetrameric channels (6-9). GIRK channels play critical roles in regulation of heart rate and aldosterone secretion (6, 7). However, subunit composition of GIRK channels in human zona glomerulosa cells of the adrenal cortex is yet to be elucidated. Notably, much lower mRNA levels of GIRK1 compared to GIRK4 have been reported in these cells (6), indicating a possible predominance of GIRK4 homotetramers.

Aldosterone, the steroid hormone secreted by the zona glomerulosa cells, is essential for blood pressure regulation. In physiological conditions, aldosterone secretion is regulated by the interplay between angiotensin II and plasma K^+^ concentration, [K^+^] (10). Angiotensin activates angiotensin II type 1 receptor, leading to depolarization of aldosterone-secreting cells through inhibition of K^+^ channels and Na^+^/K^+^ pump (11, 12). Consequently, L-and T-type voltage-gated Ca^2+^ channels (VGCC) open, Ca^2+^ enters the cell and activates aldosterone secretion. Any change in the level of angiotensin II or [K^+^] (even 1 mM) affects aldosterone secretion (10, 13).

Primary aldosteronism (PA) is a disease characterized by hypersecretion of aldosterone. PA accounts for 90% of secondary hypertension cases, approximately 10% of hypertensive patients worldwide (14). The main causes of PA are somatic and germline mutations in *KCNJ5*, *CACNA1D*, *CACNA1H*, *ATP1A1*, *ATP2B3*, *CTNNB1*, *ARMC5* genes (12). Sporadic and familial mutations in *KCNJ5* account for up to 70% of PA cases, often accompanied by adrenal adenoma (14, 15). *KCNJ5* germline mutations cause familial hyperaldosteronism type III, out of four types (16). In 2011, Choi et al. (6) showed that PA-causing mutations in the GYG motif of the selectivity filter (G151R) or in close proximity to it (T158A and L168R) in GIRK4 cause a loss of selectivity for K^+^, yielding K^+^/Na^+^ non-selective GIRK4 and GIRK1/4 channels (6). It is widely accepted that this results in depolarization caused by the influx of Na^+^ and a consequent influx of Ca^2+^ through L-and T-type VGCCs followed by constitutive aldosterone secretion and hypertension (17, 18). Since 2011, additional PA-linked mutations were discovered in *KCNJ5* gene, in the pore region or in the cytosolic N-and C-terminal domains of GIRK4 (16, 19, 20).

Loss of K^+^ selectivity and inward rectification have been reported for R52H and E246K mutations located in GIRK4 cytosolic N-and C-terminal domains, respectively (20). However, unlike for the well-established effect of mutations in the pore region (21), it is more challenging to comprehend how mutations in cytosolic domains, remote from the pore (Fig. 1), would affect selectivity. Despite that, the same gain-of-function/loss-of-selectivity mechanism suggested for PA induced by pore mutations (6) was extended for these cytosolic domain mutations as well. G247R is another mutation within the cytosolic domain found to be casually related to PA, but the mechanism remains controversial. Both loss-of-function (22) or no alteration in channel activity (20) have been reported for the G247R mutation. Thus, the biophysical and cellular mechanisms of GIRK4 channel malfunction caused by PA-linked mutations in the cytosolic domain remain incompletely understood.

**Figure 1.**
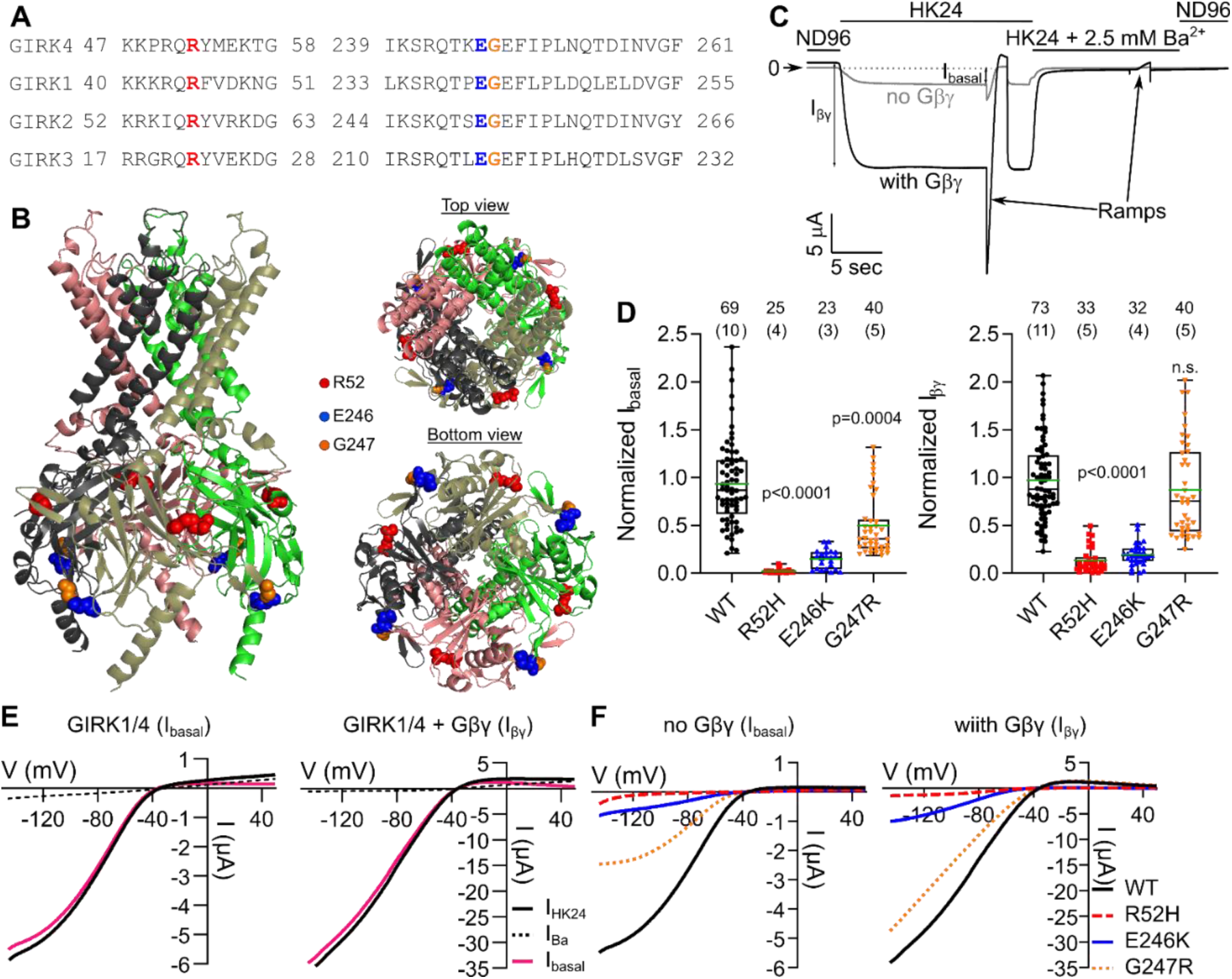
Basal and Gβγ-induced activity of heterotetrameric GIRK1/4_WT_ and mutants in *Xenopus laevis* oocytes. (A) Alignment of GIRK4 with GIRK1, GIRK2, and GIRK3 shows that R52, E246, and G247R are conserved amino acids. (B) Crystal structure of GIRK2 (23) marked with highlighted positions of the mutations; R52 – red, E246 – blue, and G247 – orange (numbering corresponds to GIRK4). (C) Example of the recording protocol in GIRK1/4_WT_ injected oocytes without (gray) or with (black) coinjected Gβγ. Horizontal bars show the corresponding external solutions. Voltage ramps were applied in HK24 and HK24 + 2.5 mM Ba^2+^ solutions. (D) Normalized I_basal_ (left) and I_βγ_ (right). Currents were measured at V_m_=-80 mV. (E) Representative I-V relationships of currents of GIRK1/4 (I_basal_; left) and GIRK1/4 + Gβγ (Iβγ_;_ right). Net GIRK currents were obtained by subtraction of I-V curve in Ba^2+^ (“Ba^2+^-subtraction” procedure). (F) Representative net I-V relationships of GIRK1/4 and mutants; I_basal_ (left) and I_βγ_ (right).

Here, we investigated GIRK4_R52H_, GIRK4_E246K_ and GIRK4_G247R_ mutants expressed in *Xenopus* oocytes and in human adrenocortical carcinoma cell line (HAC15). We demonstrate that all heterotetrameric GIRK1/4 mutants examined remain selective to K^+^ and show an unimpaired inward rectification. In contrast, the activity of heterotetrameric GIRK1/4_R52H_ and GIRK1/4_E246K_ (but not GIRK1/4_G247R_) channels is severely impaired compared to GIRK1/4_WT_. Homotetrameric GIRK4_R52H_, GIRK4_E246K_, and GIRK4_G247R_ mutants are non-functional. Single channel analysis, confocal imaging and Förster Resonance Energy Transfer (FRET) suggested both impaired gating and Gβγ interaction, and compromised plasma membrane (PM) expression in GIRK4_R52H_ and GIRK4_E246K_. For GIRK4_G247R_, the only significant defect was a severe impairment of PM expression for its homotetrameric composition. A homotetrameric GIRK4 channel opener, VU0529331, enhanced GIRK4_G247R_ currents, suggesting a potential strategy for treatment in affected patients. Our results suggest that R52H and E246K mutations are loss of function mutations that impair channel gating and membrane abundance, but not K^+^ selectivity or inward rectification. Hence, we suggest that the mechanism previously proposed for R52H and E246K mutants should be revised.

## Results

### GIRK1/4WT and GIRK1/4 mutants produce basal and Gβγ-activated currents in Xenopus laevis oocytes

The mutated amino acids (R52, E246, G247) are fully conserved in all GIRK subunits (Fig. 1A), suggesting their functional significance. They are located in the cytosolic domain far from the selectivity filter, as illustrated in Fig. 1B for the homologous amino acids in the crystal structure of GIRK2 (23).

We expressed heterotetrameric GIRK1/4 channels in *Xenopus laevis* oocytes and recorded whole-cell currents using two electrode voltage clamp. For GIRK1/4_WT_ channels, switching from low-K^+^ (2 mM K^+^) ND96 solution to a high-K^+^ (24 mM K^+^) HK24 solution yielded an inward current reflecting the GIRK’s basal activity (I_basal_). GIRK1/4_WT_ was further activated by Gβγ, which is evident from the strong increase (6.86±0.42 fold) in the constitutive K^+^ current (I_βγ_) in Gβγ-expressing oocytes (Fig. 1C). I_basal_ of GIRK1/4_R52H_ and GIRK1/4_E246K_ was significantly smaller than I_basal_ of GIRK1/4_WT_, and in some experiments it was indistinguishable from native oocyte’s currents, especially for GIRK1/4_R52H_ (Fig. 1D). To determine GIRK I-V relationships, we used voltage ramp protocols (Fig. 1E,F). The I-V curves of basal and Gβγ-evoked currents of GIRK1/4_WT_ showed the distinctive inward rectification pattern (Fig. 1E). I_basal_ of GIRK1/4_R52H_ and GIRK1/4_E246K_ often showed seemingly non-rectifying I-V relationships (Fig. 1F, left, and Fig. S1B); however, as we show below, this was an artifact arising from the very small GIRK current, comparable to oocyte’s endogenous currents. GIRK1/4_G247R_ showed distinctive inward rectification, yet I_basal_ was significantly lower compared to GIRK1/4_WT_ (Fig. 1D, F). On the other hand, I_βγ_ of GIRK1/4_G247R_ was similar to I_βγ_ of GIRK1/4_WT_, while I_βγ_ of GIRK1/4_R52H_ and GIRK1/4_E246K_ was significantly lower (Fig. 1D, F).

We also coexpressed GIRK1/4 with the muscarinic acetylcholine receptor M_2_ (M_2_R). Application of acetylcholine (ACh) evoked additional GIRK current, I_evoked_ (Fig. S1A) (24). Incidentally, however, addition of ACh also activated oocyte’s endogenous Ca^2+^ dependent chloride channels, possibly via endogenous G_q_-coupled muscarinic receptors occasionally present in the oocytes, or promiscuous coupling of m2R to G_q_ (25). These Cl^−^ currents were not sensitive to the GIRK channel blocker, Ba^2+^, decayed slowly after agonist washout, and were blocked by the injection of the Ca^2+^ chelator BAPTA (Fig. S1) (26). Ca^2+^ dependent chloride currents may be a major cause of artifact when estimating the reversal potential (V_rev_) when the expressed GIRK’s currents are small (Fig. S1B, C). To avoid these artifacts, in the following we activated the GIRK channels by co-expressing Gβγ.

### GIRK4 mutants are selective for K^+^ and retain inward rectification properties

To examine whether GIRK4 mutants lose selectivity or inward-rectification, we coexpressed GIRK1/4 channels with Gβγ and measured whole-cell currents while applying voltage ramps in four concentrations of extracellular K^+^, [K^+^]_o_ (Fig. 2A, C). V_rev_ was measured from the intercept of the net GIRK I-V curve with the voltage axis (Fig. 2B, C). Plots of V_rev_ as a function of [K^+^]_o_ (plotted on log_10_ scale, Fig. 2C) gave fairly straight lines. The slopes of the V_rev_ vs. log_10_[K^+^]_o_ plots, in mV per tenfold change in [K_o_] (decade), were 53.33±0.77, 58.49±1.6, 58.9±2.96, and 54.36±0.42 for GIRK1/4_WT_, GIRK1/4_R52H_, GIRK1/4_E246K_ and GIRK1/4_G247R_, respectively (Fig. 2D). These values are close to ~58 mV/decade predicted for a selective K^+^ channel (21). In contrast, the slope for GIRK1/4_Y152C_, a known non-selective pore mutant (27) used as control, was 13.45±4 mV/decade (Fig. 2C,D). Permeability ratios (*P*_*Na/*_*P*_*K*_) for GIRK1/4_WT_ and the R52H, E246K and G247R were close to zero, as opposed to the pore mutant GIRK4_Y152C_ (Fig. 2E). Thus, these cytosolic N-and C-terminal mutations do not affect the K^+^ selectivity. We also quantified the extent of inward rectification (F_ir_) (see Methods and Fig. 2B). F_ir_ was similar in GIRK1/4_WT_ and all mutants (Fig. 2F), usually < 0.1, except GIRK1/4_Y152C_ where F_ir_ was significantly higher, indicating partial loss of rectification in this channel, as reported (27). We conclude that mutated channels are inwardly rectifying to the same extent as WT channels.

**Figure 2.**
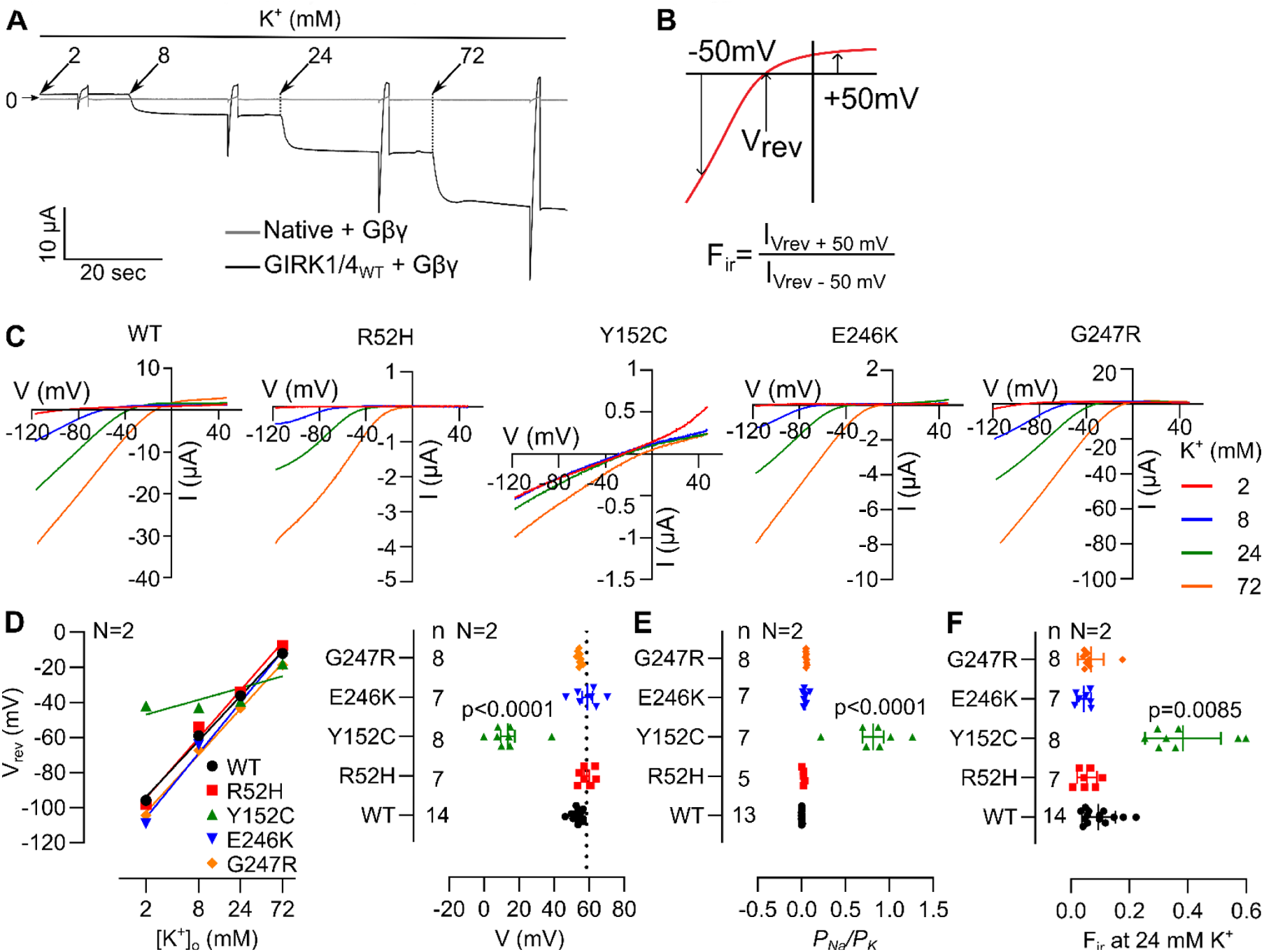
GIRK1/4_R52H_, GIRK1/4_E246K_, and GIRK1/4_G247R_ are selective for K^+^. (A) Examples of current records from oocyte expressing GIRK1/4_WT_ with Gβγ (black) and oocyte expressing only Gβγ (gray). (B) Example of F_ir_ calculation. (C) Exemplary I-V relationships of GIRK1/4_WT_ and mutants in 4 K^+^ concentrations. (D) Left panel, V_rev_ vs. log_10_[K^+^]_o_ graph of GIRK1/4_WT_ and mutants in representative oocytes. Right panel, average of slopes from the V_rev_ vs. log_10_[K^+^]_o_ graph, in mV/decade (tenfold change in [K^+^]_o_), dotted line corresponds to a slope of 58.5 mV. (E) *P_Na/_P_K_* of GIRK1/4_WT_ and mutants. (F) Factor of inward rectification (F_ir_) of GIRK1/4_WT_ and mutants.

### R52H, E246K, and G247R mutations in GIRK4 impair channels’ surface expression and interaction with Gβγ

To assess the abundance of GIRK4 mutants in the PM, we expressed GIRK4 channels tagged with YFP at the N-terminus (YFP-GIRK4), with or without Gβγ, as homo-and heterotetramers with GIRK1 (Fig. 3 and 4, respectively). Since the abundance of GIRK4 vs. GIRK1/4 in aldosterone-secreting cells remains an open question (see Introduction), we first studied the GIRK4 homotetramers (Fig. 3). Each mutant was studied in a separate experiment and compared with YFP-GIRK4_WT_. Measurement of YFP fluorescence showed a significantly lower PM expression of YFP-GIRK4_R52H_ and YFP-GIRK4_E246K_ compared to YFP-GIRK4_WT_, with or without coexpression of Gβγ (Fig. 3A, B). PM expression of YFP-GIRK4_G247R_ without Gβγ was reduced, but with Gβγ it was similar to YFP-GIRK4_WT_ (Fig 3B). Electrophysiological measurements showed that I_basal_ of GIRK4 homotetramers was very small, usually indistinguishable from native oocyte currents. Although I-V relationships occasionally showed inward rectification, the small size of GIRK currents did not allow a rigorous analysis (Fig S2). Conversely, Gβγ-evoked YFP-GIRK4_WT_ currents were quite substantial, ~ 0.5 µA on the average (Fig. 3C) and showed typical GIRK current characteristics (Fig. 3D). In contrast, I_βγ_ of all homotetrameric mutants was almost undetectable (Fig. 3C,D). This was true even for the homotetrameric YFP-GIRK4_G247R_, despite its comparable level of surface expression compared to YFP-GIRK4_WT_, in presence of Gβγ. Thus, homotetramers of the 3 mutants under study show impaired surface expression and appear non-functional.

**Figure 3.**
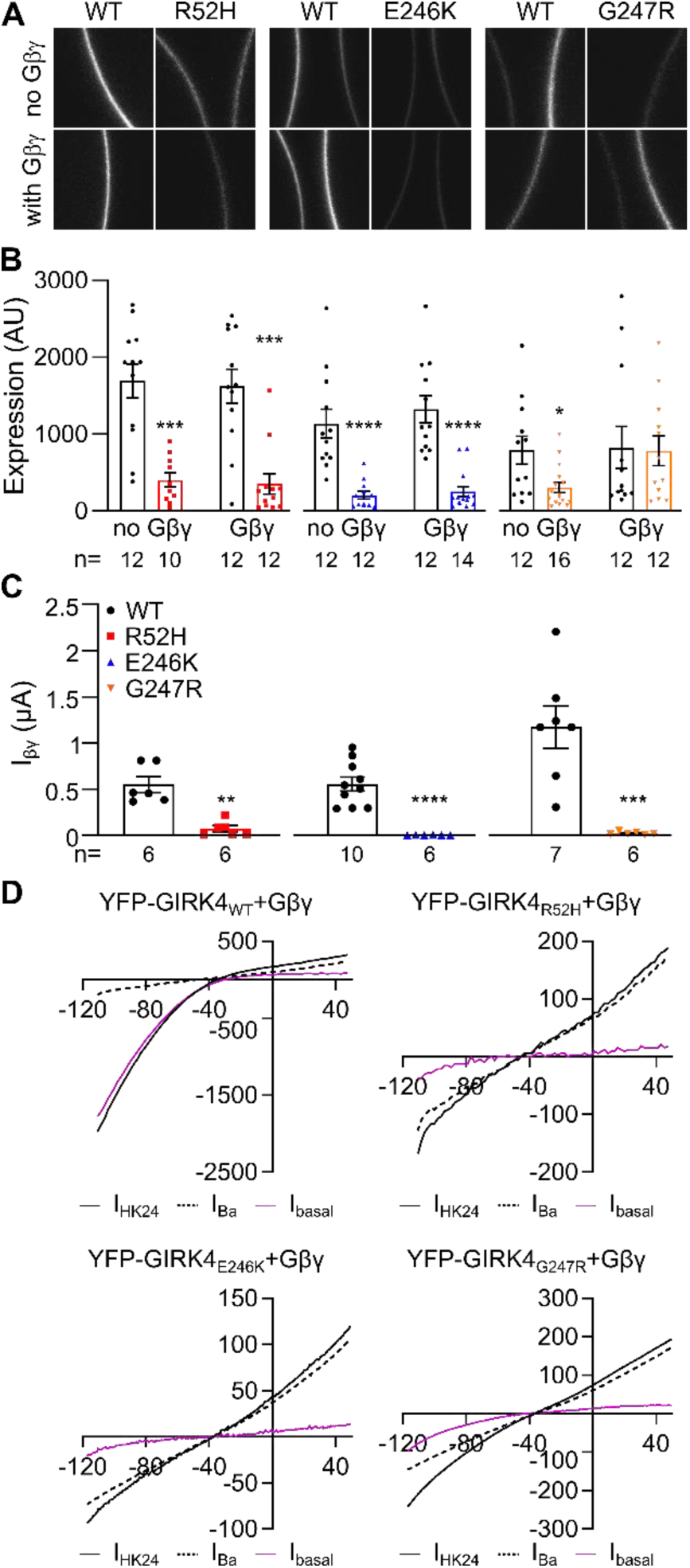
Membrane abundance of homotetrameric YFP-GIRK4 and mutants in *Xenopus laevis* oocytes. A separate experiment was performed for each mutant. (A) Representative confocal images of YFP-GIRK4 expressing oocytes, with or without Gβγ. (B) YFP-GIRK4 expression in the PM; except for YFP-GIRK4_G247R_+Gβγ, all mutants express much less than YFP-GIRK4_WT_. (C) I_βγ_ of YFP-GIRK4 homotetramers. All mutants were not active as homotetramers. (D) Representative I-V relationships of YFP-GIRK4 (WT and mutants) coexpressed with Gβγ.

**Figure 4.**
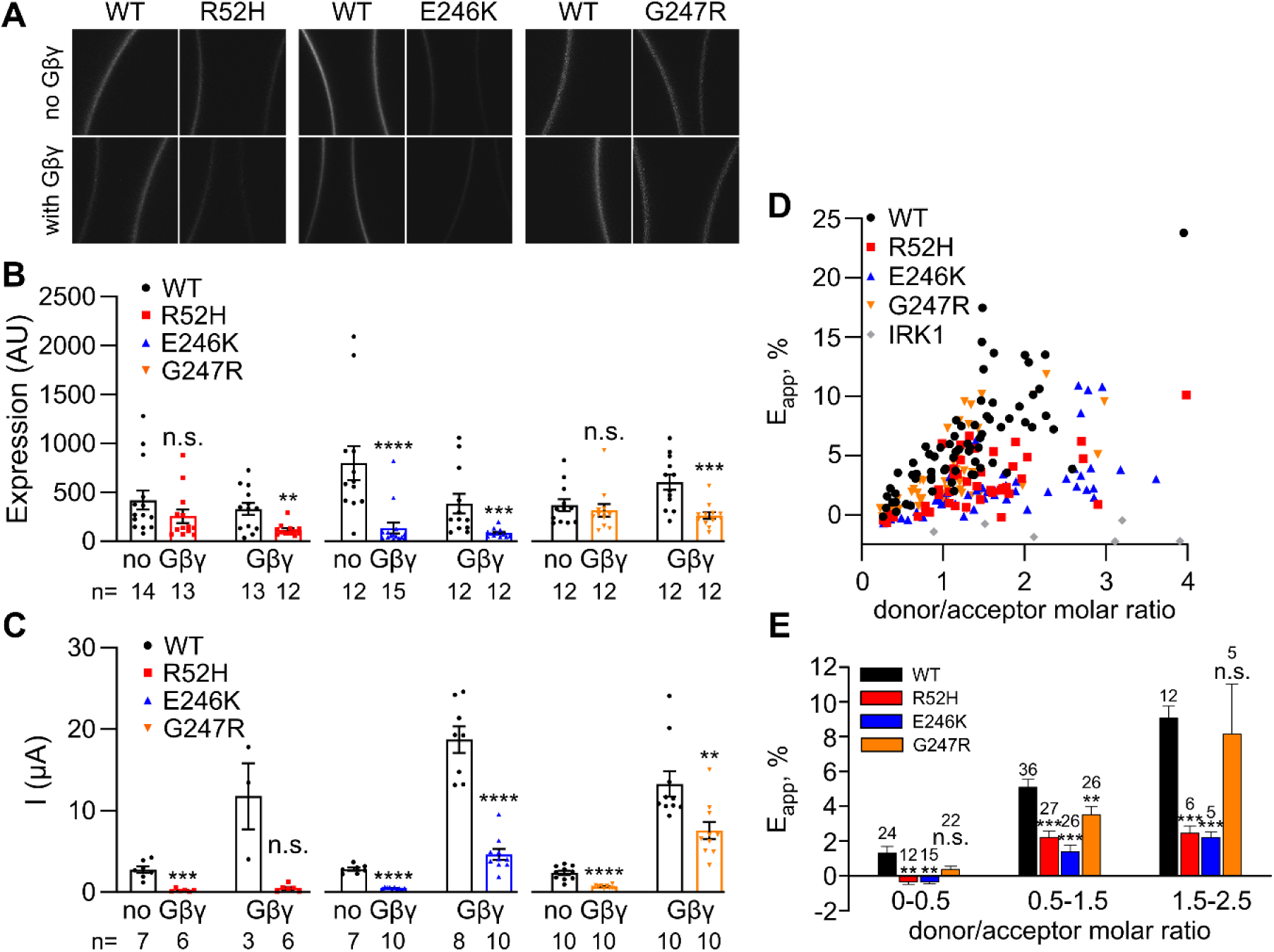
Membrane abundance of heterotetrameric GIRK1/YFP-GIRK4 and their interaction with CFP-Gβγ studied by spectral FRET. (A) Representative confocal images of GIRK1/YFP-GIRK4 expressing oocytes, with or without Gβγ. (B) GIRK1/YFP-GIRK4 expression in PM; mutated channels express less well than GIRK1/YFP-GIRK4_WT_. A separate experiment was performed for each mutant. (C) Currents of GIRK1/YFP-GIRK4_WT_ and mutated channels. (D) Raw data from FRET experiments, showing E_app_ as a function of donor/acceptor molar ratio. Each symbol corresponds to E_app_ from one oocyte, from 4 experiments. (E) Binned analysis of E_app_ shows that, at all donor/acceptor molar ratios, the interaction of CFP-Gβγ with GIRK1/YFP-GIRK4_R52H_ and GIRK1/YFP-GIRK4_E246K_ was impaired, whereas the interaction of CFP-Gβγ with GIRK1/YFP-GIRK4_G247R_ was more similar to GIRK1/YFP-GIRK4_WT_. Molar ratios >2.5 were not included in the analysis because of insufficient data points forGIRK1/YFP-GIRK4_WT_.

We next studied heterotetramers of GIRK1 with WT or mutated YFP-GIRK4. The surface expression of GIRK1/YFP-GIRK4_E246K_ was significantly lower than for the wild-type channel, with or without coexpression of Gβγ (Fig. 4A, B). The PM levels of GIRK1/YFP-GIRK4_R52H_ and GIRK1/YFP-GIRK4_G247R_ were not significantly different from GIRK1/YFP-GIRK4_WT_. However, a significant reduction for both mutants was observed when they were coexpressed with Gβγ (compared to GIRK1/YFP-GIRK4_E246K_ coexpressed with Gβγ). In the same experiments, all mutants showed reduced currents compared to WT, with or without Gβγ (Fig. 4C).

To examine whether R52H, E246K, and G247R mutations impair GIRK4 interaction with Gβγ, we used spectral FRET (28). We coexpressed CFP-Gβγ (donor; usually at two concentrations) with YFP-GIRK4 (acceptor), and GIRK1 (Fig. 4D,E, S3). YFP-labeled, G protein-insensitive inwardly rectifying K channel IRK1 (Kir2.1; *KCNJ2*) was used as negative control (Fig. S3). To circumvent variability between oocyte batches, we calculated donor/acceptor molar ratio in each oocyte using the double-labeled YFP-GIRK2-CFP (coexpressed with GIRK1) as a molecular caliper (29). In sensitized emission FRET, the FRET efficiency increases with donor/acceptor ratio (30). The FRET titration curve reaches saturation at high donor/acceptor molar ratios (30) (>5 for GIRK-Gβγ FRET pair (29)). Since in our experiments we have not achieved full saturation (Fig. 4D), we divided the data into bins with comparable donor/acceptor molar ratios, in the range where the representation of each mutant or WT was at least 3 experimental points (Fig. 4E). We found that FRET efficiency (E_app_), that reports interaction with Gβγ, was significantly impaired for GIRK1/YFP-GIRK4_R52H_ and GIRK1/YFP-GIRK4_E246K_ in all donor/acceptor molar ratio ranges. Interaction of Gβγ with GIRK1/YFP-GIRK4_G247R_ was similar or slightly smaller than with GIRK1/YFP-GIRK4_WT_ (Fig. 4E).

### Single channel analysis of GIRK4 mutants show reduced P_o_ of GIRK1/4_R52H_ but not of GIRK1/4_E246K_

To directly address the possible defects in channel function on molecular level, we measured Gβγ-induced single channel activity using cell-attached patch clamp (Figure 5A, B). We used GIRK1/4 heterotetramers, since mutated GIRK4 homotetramers were found non-functional. GIRK1/4 channels were coexpressed at low surface density, with Gβγ using doses of RNA that usually produce maximal P_o_ in GIRK1/4_WT_. Figure 5B shows exemplary recordings of GIRK1/4_WT_, GIRK1/4_R52H_ and GIRK1/4_E246K_. We found that the open probability (P_o_) of GIRK1/4_R52H_ was significantly lower than P_o_ of GIRK1/4_WT_, while P_o_ of GIRK1/4_E246K_ was not significantly different (Fig. 5C). Neither single channel current (i_single_) (Figs. 5D and S4B) nor single channel conductance (g) (Figs. 5E and S4C) were affected by the mutations.

**Figure 5.**
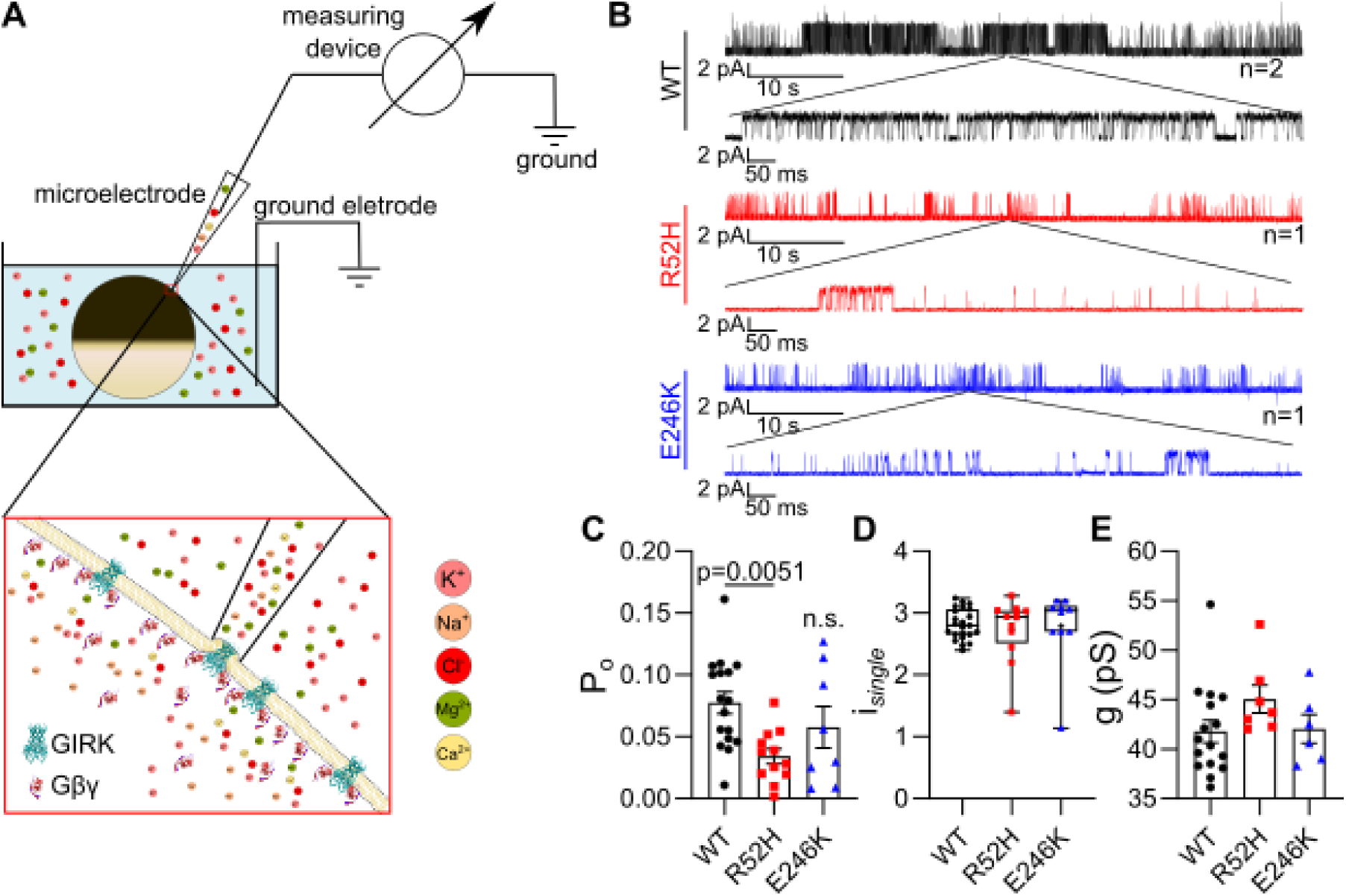
Cell attached patch clamp recording and single channel analysis of heterotetrameric GIRK1/4 channels. (A) A simplified scheme of the cell attached patch clamp recording. (B) Representative recordings of GIRK1/4_WT_, GIRK1/4_R52H_, and GIRK1/4_E246K_ with a high dose of Gβγ (5 ng Gβ and 1 ng Gγ RNA). (C) P_o_ of GIRK1/4_R52H_ is significantly reduced compared to GIRK1/4_WT_. (D) i*_single_* of mutants and (E) single channel conductance of mutants are similar to GIRK1/4_WT_.

Another way to assess mutation-induced changes in P_o_ is from whole-cell data of Figs. 3 and 4. The whole-cell current, I, is given by the equation I=i_single_*P_o_\**N* (21), where *N* is the total number of channels in the PM. *N* is proportional to fluorescence intensity of the labeled channels (in arbitrary units), E_AU_. Thus, whole-cell current normalized to PM level, I/E_AU_, is proportional to I/*N*, therefore I/E_AU_ ~ i_single_*P_o_. Since i_single_ is not affected by mutations, differences in I/E_AU_ reflect differences in P_o_. The homotetrameric YFP-GIRK4 mutants exhibited a greatly reduced I/E_AU_ (Fig. S5A), suggesting a great reduction in P_o_ (impaired gating) in addition to the impaired expression. For heterotetrameric GIRK1/YFP-GIRK4 mutants, the normalization procedure showed an impaired P_o_ for GIRK1/YFP-GIRK4_R52H_ but not for GIRK1/YFP-GIRK4_E246K_ or GIRK1/YFP-GIRK4_G247R_, supporting the single channel results and indicating that the E246K and G247R mutations do not impair the gating properties of heterotetrameric GIRK1/4 channels (Fig. S5B).

### Expression of GIRK1/4 in HAC15 cells

The type of K^+^ channels in the zona glomerulosa cells varies between species, GIRK4 is not present in the adrenal cortex of rats, and probably also in mice (31). Thus, rodent models are ineffective to study the effect of GIRK4 mutations in vivo (31, 32). Hence, to corroborate that mutation-induced changes in GIRK4 function observed in *Xenopus* oocytes also take place in a more relevant cellular environment, we used the human adrenocortical carcinoma cell line (HAC15). First, we tested the presence of endogenous GIRK1 and GIRK4 using Western blot. Both GIRK1 and GIRK4 proteins were present in these cells (Fig. 6A). However, whole-cell recordings of untransfected cells, or cells transfected with D_2_ dopamine receptor, did not show any basal or dopamine (DA)-evoked GIRK current (Fig. 6B, left). Next, we transfected the cells with GFP-GIRK1 and GIRK4 (WT or mutants), with Gβγ subunits and the D_2_ receptor. I_evoked_ was elicited by application of 100 µM dopamine. The activity of GIRK1/4_R52H_ and GIRK1/4_E246K_ mutants was significantly reduced compared to GIRK1/4_WT_, while GIRK1/4_G247R_ did not differ from GIRK1/4_WT_ (Fig. 6B).

**Figure 6.**
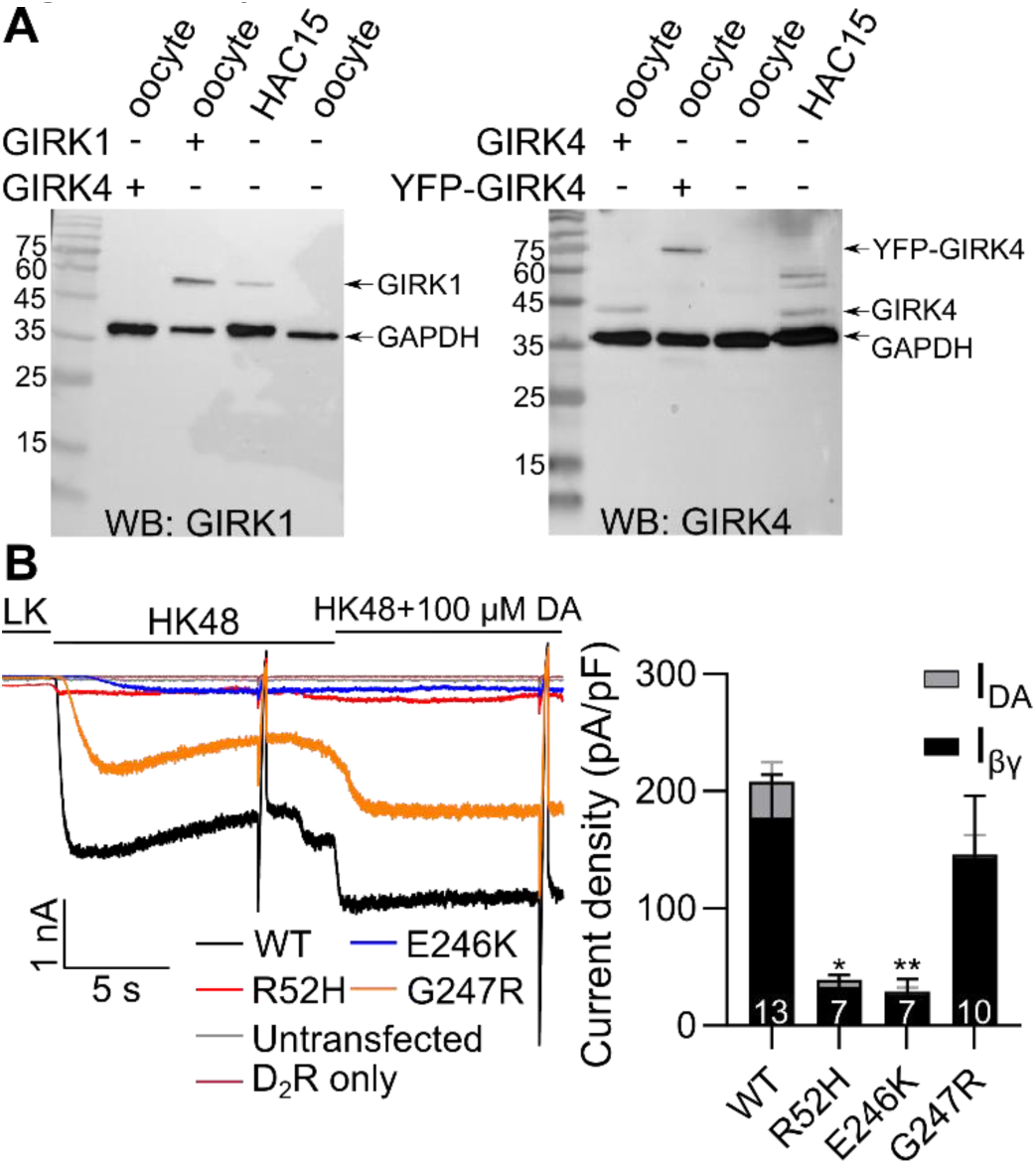
Expression of GIRK4 mutants in HAC15 cells. (A) Western blots of HAC15 cells with GAPDH antibody (both panels) and GIRK1 antibody (left) and GIRK4 antibody (right) show that HAC15 cells express GIRK1 and GIRK4 channels. Oocytes expressing GIRK1, GIRK4, and YFP-GIRK4 were used as positive controls, naïve oocytes as negative control, and GAPDH as loading control. (B) Examples (left) and summary (right) of currents in HAC15 cells expressing GFP-GIRK1/GIRK4_WT_, GFP-GIRK1/GIRK4_R52H_, GFP-GIRK1/GIRK4_E246K_, and GFP-GIRK1/GIRK4_G247R_. Channels were expressed with D_2_ dopamine receptor and Gβγ (I_βγ_), thus the small I_DA_.

### The GIRK2 and GIRK4 opener VU0529331 activates GIRK4_G247R_ homotetramer

Lastly, we aimed to test whether the activity of mutated channels can be rescued by the recently discovered GIRK2 and GIRK4 opener VU0529331, which activates the channels in a Gβγ-independent manner (33). We used oocytes expressing GIRK4 homotetramers, with or without Gβγ. VU0529331 activated GIRK4_WT_ and GIRK4_G247R_ in a dose-dependent manner, and the activated channels showed the typical inwardly rectifying I-V relationships (Fig. 7). No response could be seen in oocytes injected with GIRK4_R52H_ and GIRK4_E246K_ RNA.

**Figure 7.**
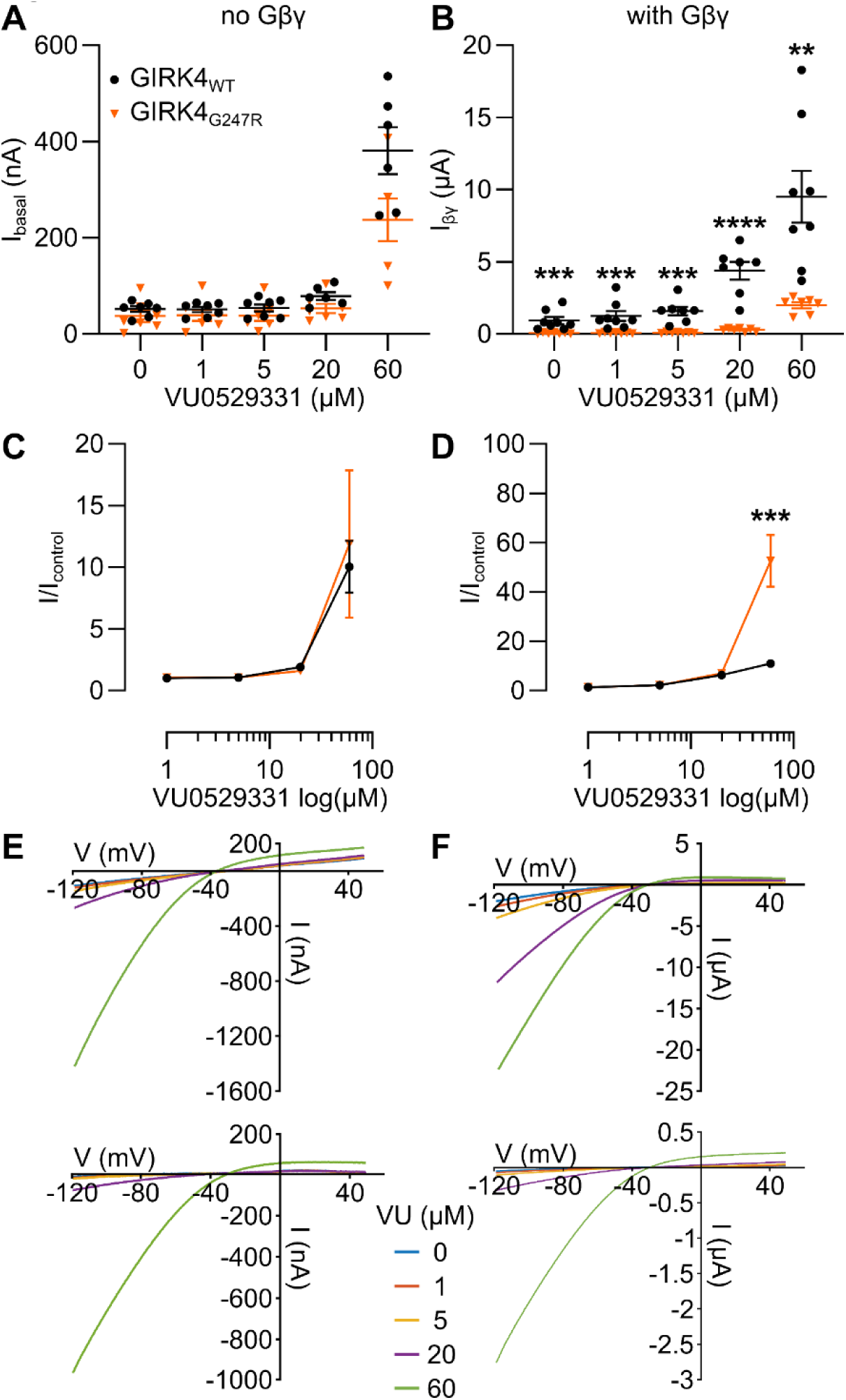
The effect of VU0529331 on homotetrameric GIRK4 mutants. Oocytes expressed GIRK4 mutants and GIRK4_WT_, with or without Gβγ. Cells were exposed to 4 concentrations of VU0529331 (1, 5, 20, and 60 μM). Only GIRK4_WT_ and GIRK4_G247R_ mutant responded to VU0529331. (A) I_basal_ of GIRK4_WT_ and GIRK4_G247R_. GIRK4_WT_ and GIRK4_G247R_ react similarly to VU0529331. (B) I_βγ_ of GIRK4_WT_ and GIRK4_G247R_. GIRK4_WT_ had larger current at 0 μM VU0529931, while GIRK4_G247R_ did not show any response to concentrations lower than 20 μM VU0529331. (C) Normalized I_basal_ (I/I_control_). GIRK4_WT_ and GIRK4_G247R_ were activated similarly by VU0529331. (D) I/I_control_ of I_βγ_ show that GIRK4_G247R_ was activated by VU0529331 stronger than GIRK4_WT_, in relative terms. (E) I-V relationship of GIRK4_WT_ (top) and GIRK4_G247R_ (bottom) without Gβγ. (F) I-V relationship of GIRK4_WT_ (top) and GIRK4_G247R_ (bottom) with Gβγ.

## Discussion

Loss of selectivity of GIRK4 (*KCNJ5*) channels is believed to be the leading cause of PA, by producing constitutive depolarization of zona glomerulosa cells (6, 17, 27). This concept was extended from mutations in the GIRK4 pore region (6, 27, 34) to additional mutations in the transmembrane domain of GIRK4 (34, 35) and, eventually, to mutations in the cytosolic N-and C-termini (17, 20), despite the identification of two cytosolic domain mutations (R115W and E246G) that impaired membrane abundance of GIRK4 rather than selectivity (19). We studied PA-linked GIRK4 mutations of three conserved amino acids located in the cytosolic domain (Fig. 1): R52 and E246, that were reported to cause loss of selectivity of GIRK4 channels (20), and G247. Here, we confront the selectivity loss concept for C-and N-terminal mutations; we show that R52H, E246K and (to a milder extent) G247R mutations cause a complete or partial loss of function and/or channel abundance in PM, but do not alter channel selectivity or inward rectification.

Many of the *KCNJ5* mutant studies (including our work) were done in *Xenopus laevis* oocytes used as heterologous expression system (20, 22, 34, 36-38). We first thoroughly investigated possible sources of artifact in this system. Like other cells, *Xenopus* oocytes possess a number of endogenous ion channels that yield small currents, usually in the range of a few tens of nA, but may give rise to artefactual interpretations if the expressed exogenous ion channels show poor expression or function. For instance, activation of M2 receptors by ACh often elicited outwardly-rectifying endogenous Ca^2+^-dependent Cl^−^ currents (25) that were abolished by the Ca^2+^ chelator BAPTA (Fig. S1). Even the remaining leak currents may be mistaken for currents of GIRK mutants if the expressed currents are small. To avoid these artifacts, we activated GIRK channels by coexpressing Gβγ subunits at doses that maximally activate the WT GIRKs; yet, even under these conditions the homotetrameric GIRK4_R52H_ and GIRK4_E246K_ did not produce detectable currents.

Another source of ambiguity in GIRK4 mutant studies is the subunit composition. In view of the low abundance of GIRK1 RNA in zona glomerulosa (6), it is conceivable that GIRK4 homotetramers are abundant in aldosterone-secreting cells. However, most heterologous expression studies have been done with GIRK1/4 heterotetramers (19, 27, 35, 36, 39). Only two GIRK4 pore mutants have been tested as homotetramers, and reported to be functional and non-selective (6). Homotetrameric GIRK4_G247R_ mutants was tested on the background of an additional mutation, S143T (22). The homotetrameric GIRK4_S143T_ is a pore mutant that expresses better than GIRK4_WT_ and preserves inward rectification (40, 41), but its selectivity has not been carefully characterized. The GIRK4_S143T_ homotetramers showed large I_basal_ (22), which is different from GIRK4_WT_ homotetramers that have very small I_basal_ and large I_βγ_, as we have demonstrated (Fig. S2). To address this conundrum, we examined both homotetrameric and heterotetrameric (with GIRK1) GIRK4, WT and mutants.

Our central finding was that, in GIRK1/4 heterotetrameric context, all mutations strongly reduced the whole-cell basal GIRK current in *Xenopus* oocytes, and R52H and E246K (but not G247R) mutations greatly reduced the Gβγ-induced currents, both in in *Xenopus* oocytes and in HAC15 cells (Figs. 1 and 6). The homotetrameric mutant channels gave practically no detectable currents at all (Fig. 4). These results indicate that all three mutations cause loss of function, which is more severe in homotetrameric than in heterotetrameric channels.

In contrast, we found no changes in K^+^ selectivity or inward rectification (in the heterotetrameric context; Fig. 2). The transmembrane pore of K^+^ channels is a conserved structure that enables K^+^ ions to permeate rapidly through the channel over Na^+^ ions, and mutations in the pore region shift the permeability ratio towards Na^+^ (42, 43). The cytosolic domain of inwardly rectifying K^+^ channels forms an additional, cytoplasmic pore that is a continuation of the transmembrane pore but does not contain a selectivity filter (44-46). Several amino acids of the cytosolic domain lining the pore control channel’s inward rectification by altering Mg^2+^ and polyamine block, but not the ion selectivity (5, 46, 47). We measured the reversal potential of Gβγ-activated GIRK1/4 current in four external K^+^ concentrations and found a linear dependence between log[K^+^] and V_rev_ with a slope close to that predicted by the Nernst equation for a pure K^+^ pore, and identical to that of the wild type channel. Analysis using the Goldman–Hodgkin–Katz equation confirmed that the mutated channels conducted K^+^ but not Na^+^. Evaluation of the degree of inward rectification showed that inward-rectification of mutated channels was similar to WT (48). Our results clearly demonstrate that all mutants studied here are selective to K^+^ over Na^+^, and rectify, to a similar extent.

To understand the mechanisms underlying loss of function of the mutants, we employed several approaches. Confocal imaging in *Xenopus* oocytes revealed that PM abundance of mutated homo- or heterotetrameric channels was significantly reduced. Analysis of macroscopic whole-cell currents normalized to PM expression indicated impaired gating (in addition to reduced PM abundance) of R52H, and impaired PM abundance, but not gating, of E246K and G247R heterotetramers (Fig. S5). Single channel recordings (Fig. 5) confirmed the reduction in open channel probability (P_o_), which indicates impaired gating, of the heterotetrameric GIRK1/4_R52H_. Finally, FRET experiments suggested an impaired interaction of the R52H and E246K mutant channels with the main gating factor, Gβγ (Fig. 4D,E). On the other hand, while the heterotetrameric GIRK1/4_E246K_ had an impaired interaction with Gβγ (FRET data), we did not see significant reduction in P_o_. Hence, we suggest that R52H mutation impairs both gating and PM abundance of homo- and heterotetrameric channels. E246K mutation impairs mainly channel expression, and to a lesser extent channel activity (in the heterotetrameric context), mainly because of the reduced Gβγ affinity. G247R mutation had the smallest impact on the heterotetrameric channel, and showed mildi mpairment in activity (I_basal_, but not I_βγ_) and in PM abundance.

In our study, none of the mutated homotetramers produced detectable whole-cell currents, with or without coexpression of Gβγ. Homotetrameric GIRK4_G247R_ showed PM expression similar to the WT homotetramer, but it was not activated by Gβγ. Remarkably, however, it was activated by the GIRK4 opener, VU0529331. Since the heterotetrameric GIRK1/4_G247R_ does not show a significant loss of function, yet the patients with G247R mutation express a pathologic phenotype, these results indicate that the subunit composition in human aldosterone-secreting cells tends toward homotetrameric GIRK4 channels rather than heterotetrameric GIRK1/4.

On the other hand, GIRK4_R52H_ and GIRK4_E246K_ showed very low expression in the PM compared to GIRK4_WT_, and this could be the main reason for the absence of current. These two mutants could not be activated by VU0529331 or Gβγ. However, since we see some expression, it is possible that other openers will be able to activate these channels.

### Conclusions and prospects

Taken together, our results suggest that the mechanism proposed for R52H and E246K mutants (20) should be revised. We propose that, as opposed to pore mutations, N-and C-terminal domain mutations studied here reduce channel activity and PM abundance. This results in a decrease in resting K^+^ conductance, therefore the resting membrane potential becomes more positive, which triggers Ca^2+^ flow into the cell and hypersecretion of aldosterone. These results are supported by the previous study of Cheng et al. (19), which showed reduced activity and PM abundance of two additional N-and C-terminal domain mutants. Knowing the exact biophysical mechanism that impairs the channel is crucial for setting the course of treatment. While patients with pore mutations, which yield gain-of-function non-selective Na^+^/K^+^ channels, may benefit from treatment with channel blockers, patients with cytosolic domain mutations may potentially benefit from treatment with GIRK4 channel openers to achieve the same effect: prevention of constitutive depolarization of zona glomerulosa secretory cells.

## Materials and Methods

### DNA constructs and RNA

cDNA constructs of rat GIRK1, N-terminally YFP-tagged GIRK1 (YFP-GIRK1), GIRK4_WT_, GIRK4_R52H_, GIRK4_E246K_, and GIRK4_G247R_ were inserted into pBS-MXT vector. DNA of human N-terminally YFP-tagged GIRK4_WT_ (YFP-GIRK4_WT_) was a gift from Prof. Wolfgang Schreibmayer, and mutations were made to produce YFP-GIRK4_R52H_, YFP-GIRK4_E246K_, and YFP-GIRK4_G247R_. mRNAs were prepared as described previously (24).

The amounts of mRNA injected per oocyte were varied according to the experimental design, in ng/oocyte: 0.5-2 GIRK1, 0.5-2 YFP-GIRK1, 0.25-1 GIRK4, 0.25-5 YFP-GIRK4. For maximal channel activation by Gβγ, we injected 5 ng Gβ and 1-2 ng Gγ mRNA. These weight ratios were chosen to keep approximately equal molar amounts of Gβ and Gγ RNA. RNA of the muscarinic 2 receptor (M_2_R), if present, was 1 ng. 25 ng of the anti-GIRK5 oligo nucleotide antisense XIR was routinely injected to prevent the formation of GIRK1/5 channels (49).

### Ethical approval of *Xenopus laevis*, oocyte preparation, and electrophysiology

Experiments have been approved by Tel Aviv University Institutional Animal Care and Use Committee (permit #01-16-104). Maintenance and surgery of female frogs were as described previously (50). All materials were from Sigma unless indicated otherwise. Oocytes were defolliculated by collagenase, injected with RNA and incubated for 3 days at 20–22^◦^C in ND96 solution (low K^+^) (in mM: 96 NaCl, 2 KCl, 1 MgCl_2_, 1 CaCl_2_, 5 HEPES, pH 7.5), supplemented with 2.5 mM sodium pyruvate, 100 µgml^−1^streptomycin and 62.75 µgml^−1^ penicillin (or 50 µgml^−1^ gentamycin) (24). Whole-cell GIRK currents were measured using standard two-electrode voltage clamp method at 20–22^◦^C, in different K^+^ concentrations (8, 24, 48, 72 or 96 mM K^+^ (HK)) (51). Different concentrations of K^+^ solutions were obtained by mixing ND96 with a 96 mM K^+^ solution containing, in mM: 96 KCl, 2 NaCl, 1 CaCl_2_, 1 MgCl_2_, 5 HEPES, pH adjusted to 7.5 with KOH. Acetylcholine (ACh) was added to HK solution as agonist at 10 µM, Ba^2+^ (2.5 mM) was used to block the channel. Current-voltage (I-V) relations were obtained using 2 s voltage ramps from −120 mV to +50 mV. Net GIRK I-V relationships were obtained by subtracting the current remaining after blocking all GIRK activity with 2.5 mM Ba^2+^. In one series of experiments (Fig. 2), we obtained the net GIRK I-V relationship by subtracting averaged I-V relationship from native oocytes injected with only Gβγ. For small GIRK currents (in low [K^+^]_out_) this procedure is superior over subtraction of Ba^2+^-blocked currents, avoiding possible inaccuracies caused by different extent of Ba^2+^ block of inward and outward GIRK currents(51).

VU0529331 (Alomone Labs; V-155), an opener of homotetrameric GIRK2 and GIRK4 channels, was dissolved in 100% DMSO to a final concentration of 25 mM. To measure GIRK4 response to VU0529331, the drug was diluted into HK24 solution to get HK24 + 60 µM VU0529331 solution. Consecutively, this solution was diluted again to get 20 µM, and so on for all concentrations.

### HAC15 cells culture, transfection, biochemistry and electrophysiology

Cells were acquired from ATCC (ATCC® CRL3301 ™) and cultured as described (52). Cells were grown in Dulbecco’s Modified Eagle’s/Ham’s F-12 medium (DMEM/F-12) (Gibco #11330-05) containing 10% Cosmic calf serum (Hyclone #SH30), 1% L-Glutamine (Sigma-Aldrich; A7506), 1% ITS (Becton Dickenson - FAL354352) and penicillin– streptomycin (Sigma-Aldrich; P4333).

For Western blot, cultured HAC15 cells were treated as described previously (53). GIRK1/4 RNA-injected oocytes were used as positive control. Oocytes were homogenized on ice in homogenization buffer (20 mM Tris, pH 7.4, 5 mM EGTA, 5 mM EDTA, and 100 mM NaCl) containing protease inhibitor mixture (Roche Applied Science). The homogenates were centrifuged at 3000 × *g* for 5 min at 4 °C and the pellet was discarded. Protein samples (35µg) were electrophoresed on 12% polyacrylamide-SDS gel and transferred to nitrocellulose membranes for Western blotting with antibodies; anti-GIRK1 at 1:300 dilution (Alomone Labs, APC-005) or anti-GIRK4 at 1:100 dilution (Alomone Labs, APC-027), and GAPDH in 1:1000 or 1:2000 dilution (Cell Signaling Technology, P04406). Goat Anti-Rabbit IgG Antibody, (H+L) HRP conjugated secondary antibody at 1:40,000 dilution was applied (Jackson ImmunoResearch Labs 111-035-144, RRID:AB_2307391).

For electrophysiological experiments, cells were plated in a 6-well plate and were transfected with combinations of the following DNAs as designed for specific experiments: 0.5 ng GFP-GIRK1, 0.5 ng GIRK4_WT_, 0.5 ng GIRK4_R52H_, 0.5 ng GIRK4_E246K_, 0.5 ng GIRK4_G247R_, 0.5-1 ng Gβ_1_, 0.5-1 ng Gγ_2_ (all in pcDNA3.1); 0.5 ng D_2L_ receptor in pXOOM (the latter was kindly provided by Kristoffer Sahlholm). 24-36 hours after transfection cells were moved onto 13 mm cover slips, coated with fibronectin and electrophysiological measurements were performed in the next 24-48 hours.

Whole-cell currents were measured using Axopatch 200B (Molecular Devices, Sunnyvale, CA). Holding potential was −80 mV, and voltage ramps were from −120 to +20 mV, no correction for junction potential (~13 mV) was made. Currents were recorded in low K^+^ solution (in mM): NaCl 136, KCl 4, CaCl_2_ 2, MgCl_2_ 2, HEPES 10, NaH_2_PO_4_ 0.33, Glucose 10, pH 7.4, or in high K^+^ (HK48) solution (in mM): NaCl 92, KCl 48, CaCl_2_ 2, MgCl_2_ 2, HEPES 10, NaH_2_PO_4_ 0.33, Glucose 10, pH 7.4. 100 μM dopamine was used to measure GPCR response. Electrode solution contained (in mM): NaCl 6, KCl 22, K-Gluconate 110, MgCl_2_ 2, HEPES 10, EGTA-HOH 1, ATP-K_2_ 2, GTP-Tris 0.5, pH 7.2.

### Cell-attached single channel recordings in *Xenopus* oocytes

Patch clamp experiments were done using Axopatch 200B as described (54). Currents were recorded at −80 mV, filtered at 2 kHz and sampled at 20 kHz. Patch pipettes had resistances of 1.4–3.5 MΩ. Pipette solution contained, in mM: 146 KCl, 2 NaCl, 1 MgCl_2_, 1 CaCl_2_, 1 GdCl_3_, 10 HEPES/KOH (pH 7.5). GdCl_3_ completely inhibited the stretch-activated channels. The bath solution contained, in mM: 146 KCl, 2 MgCl_2_, 1 EGTA, 10 HEPES/KOH (pH 7.5). To obtain single channel recordings, oocytes were injected with low doses of RNA of GIRK1 (10–100 pg/oocyte), and RNA of GIRK4 was half of that (5-50 pg); the amount of GIRK1/4_WT_ RNA was 10-20/5-10 pg, respectively. The amounts of the GIRK1/4_R52H_ and GIRK1/4_E246K_ channels’ RNA were 20-40/10-20 pg and 40-100/20-50 pg, respectively. In addition, 25 ng of the antisense DNA oligonucleotide against the endogenous GIRK5 channel was injected together with the RNAs (49). Single channel current (i*_single_*) was calculated from all-point histograms of the original records (55), and open probability (P_o_) was obtained from event lists generated using idealization procedure based on 50% crossing criterion (56). Number of channels was estimated from overlaps of openings during the whole time of recording (at least 5 min). P_o_ was calculated only from records that contained up to 3 channels. Thus, the probability of missing a channel was negligible. For channels activated by co-expressed Gβγ there was no decrease in P_o_ over >4 minutes, and the P_o_ was averaged from the first 4 minutes of the record. Single-channel conductance (g) was measured from I-V relationships, constructed from values of i_single_ measured at potentials ranging from −120 mV to −40 mV at 20 mV increments.

### Confocal imaging

Confocal imaging of the oocytes and analysis were performed as described (29), with a Zeiss 510 META confocal microscope, using a 20× objective. In whole oocytes, the image was focused on an oocyte’s animal hemisphere, at the equator. Images were acquired using spectral (λ)-mode. YFP was excited with the 514 nm line of the argon laser and sampled at 534–546 nm. Fluorescence signals were averaged from three regions of interest using Zeiss LSM Image Browser, background and the average signal from uninjected oocytes were subtracted.

### Förster resonance energy transfer (FRET)

FRET experiments were performed using sensitized emission spectral method (28) as described previously (29) (see also Supplemental Fig. S3). Two spectra were collected from the animal hemisphere of each oocyte, with 405nm (CFP excitation) and 514nm (YFP excitation) laser lines. Oocytes with low signal of one of the fluorophores (usually < 100 AU), or with coefficient of variation of A parameter at 524, 535, 545, 566 nm of more than 0.25, were excluded from analysis. Net FRET signal was calculated in the YFP emission range (with the 405 nm excitation) by consecutive subtraction of a scaled CFP-only spectrum (giving the A ratio parameter) and then of the ratio A0, which reports the direct excitation of YFP by the 405 nm laser, as in Equations 1 and 2:

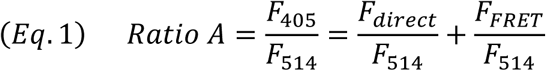

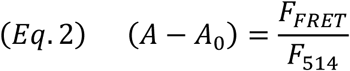

Because of the use of different experimental settings in independent experiments, the fluorescence intensities or A ratios cannot be used to construct FRET titration curves. We imaged a double-labeled protein, expressing both CFP and YFP at a 1:1 stoichiometry (YFP-GIRK2-CFP, DL-GIRK2) in each experiment to convert the fluorescence of CFP and YFP into their molar ratio (the average donor/acceptor ratio, in arbitrary units (AU) corresponds to a molar ratio of one) (29). The molar ratio collected and from many experiments can then be implemented into one titration curve (29).

The apparent FRET efficiency in an individual cell, E_app_, was calculated as in Equation 3:

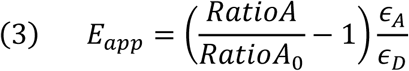

where *∈*_*D*_ and *∈*_*A*_ are molar extinction coefficients for the donor and acceptor, respectively, at the donor excitation wavelength (29, 30).

### Data analysis

I-V curves were analyzed using scripts written in-house and from Mathworks forum for analyzing. abf files produced by Clampex (Molecular Devices), MATLAB R2015a (MathWorks, USA), and with Clampfit 10.7 software (Molecular Devices). The reversal potential (V_rev_) was determined from the intercept of the net I-V curve with voltage axis.^24,27^ The extent of inward rectification (F_ir_) was determined by dividing the current at 50 mV positive to V_rev_ by the current at 50 mV negative to V_rev_ (see Fig. 2B) (48):

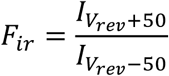

Estimated reversal potential (V_rev_) was calculated using the Nernst equation:

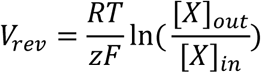

Permeability ratio of sodium and potassium (pNa^+^/pK^+^) was determined from Goldman– Hodgkin–Katz equation:

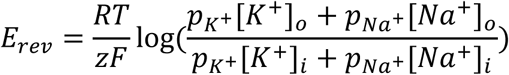

### Statistical analysis

Protein expression imaging data have been normalized as described previously (57). For analysis across several experiments, fluorescence intensity in each oocyte was normalized to the average signal in the oocytes of the control group of the same experiment. This procedure yields average normalized intensity as well statistical variability (e.g. SEM) in all treatment groups as well as in the control group. Statistical analysis was performed using SigmaPlot 11 (Systat Software, Inc.) and GraphPad Prism version 8 for Windows (GraphPad Software, La Jolla California USA). If the data passed the Shapiro–Wilk normality test and the equal variance test, two-group comparisons were performed using t-test, while multiple group comparison was performed using one-way ANOVA. If not, the Mann–Whitney rank sum test, or Kruskal-Wallis test were performed, respectively. The data in the bar graphs are presented as mean ± SEM. In box plots, boxes show 25^th^ and 75^th^ percentiles and whiskers show minimal and maximal values, black horizontal line – median, green line -mean.

## Acknowledgments

We are grateful to Alomone Labs for the generous gift of VU0529331 and to Mariam Ashkar for assistance in some of the experiments. This work was supported by the joint Israel-India grant ISF # 2255/15 and the Israel Science Foundation grant # 1282/18.

## Supplementary figures

**Figure S1.**
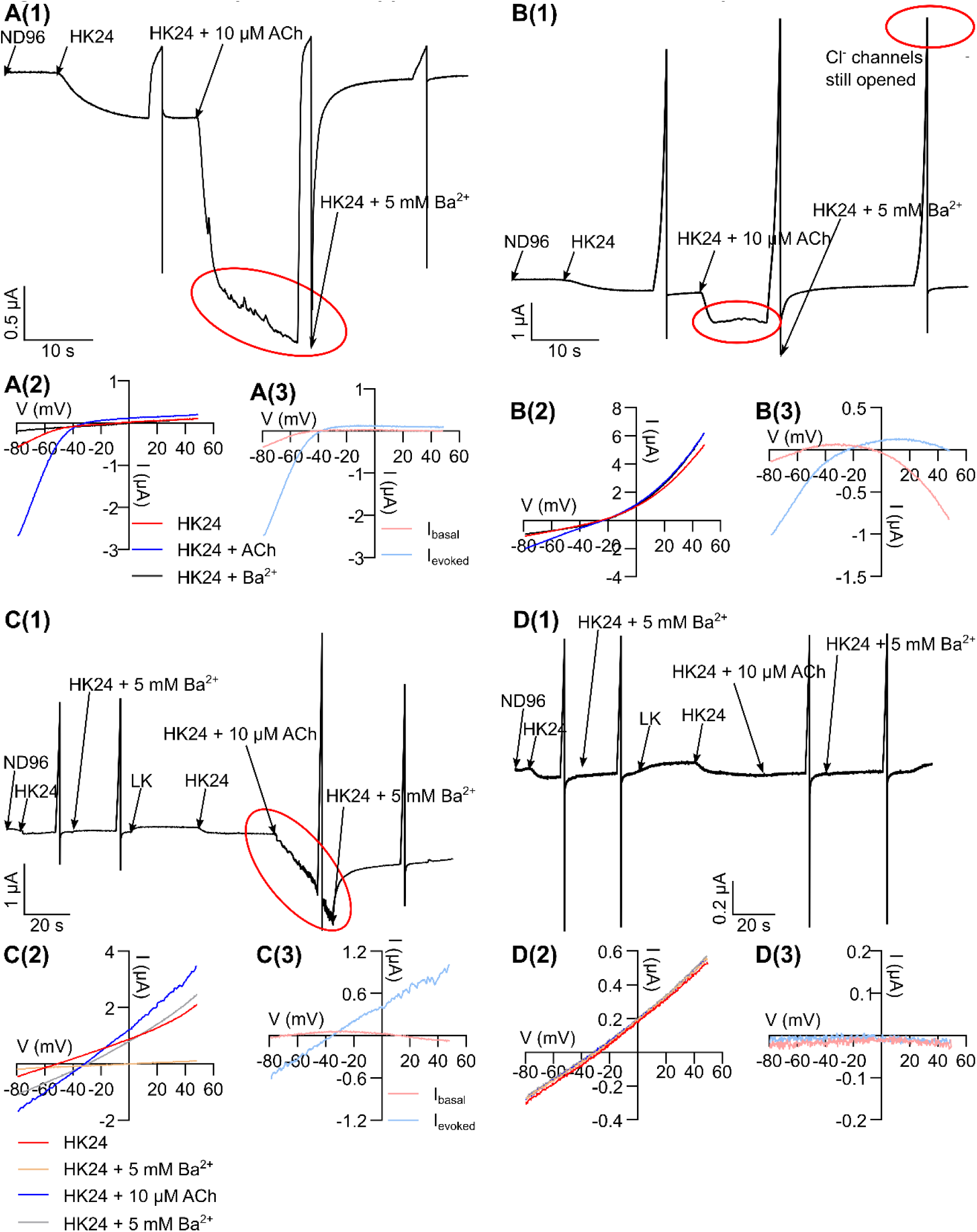
In some oocyte batches, application of ACh induces Ca^2+^-dependent Cl^−^ currents. (A) ACh evokes Ca^2+^-dependent Cl^−^ currents in oocytes expressing GIRK1/4 and M2R. (A1) Example of a recording from an oocyte coinjected with RNAs of GIRK1/4_WT_ and M_2_R. Application of ACh evoked GIRK’s I_evoked_ but also a Ca^2+^-dependent Cl^−^ current (marked with red circle). (A2) I-V relationships for I_basal_ and I_evoked_ before Ba^2+^ subtraction. (A3) Net I-V relationships of I_basal_ and I_evoked_ after Ba^2+^ subtraction. Cl^−^ currents were masked by the large currents of GIRK1/4_WT_. (B) ACh-induced Cl^−^ currents significantly contribute to the overall small currents in an oocyte expressing GIRK1/4R52H and m2R. (B1) The full current record. Application of ACh evoked a Ca^2+^ dependent Cl^−^ current (marked with red circle), which was not blocked by Ba^2+^ and decayed slowly, over minutes. Since the currents of GIRK1/4R52H were small, the Cl^−^ current contributed a large part of the total current. (B2) I-V relationship before Ba^2+^ subtraction. (B3) I-V relationship of I_basal_ and I_evoked_ after Ba^2+^ subtraction. Because Cl^−^ currents (which are outwardly rectifying and thus larger at positive voltages) are still present in Ba^2+^ after ACh washout, the subtraction of I-V curve recorded in the presence of Ba^2+^ results in an I-V curve showing inward currents at positive voltages, which is an artifact. (C) ACh may evoke Cl^−^ currents in naïve (uninjected with RNA) oocytes. (C1) Example of a recording from an uninjected oocyte; here, Ba^2+^ was applied after HK24, then washed with ND96, then HK24 was applied again followed by ACh, and ACh was washed out with ND96 + 5 mM Ba^2.^. This protocol was developed so we could get net I_basal_ and I_evoked_, and yet, the artifacts remained. (C2) I-V relationship before Ba^2+^ subtraction. (C3) I-V relationship of Ba^2+^ subtracted I_basal_ and I_evoked_. (D) BAPTA eliminated the Ca^2+^ dependent Cl^−^ current. (D1) Example of a recording from an uninjected oocyte ~1 hour after the injection of BAPTA. (D2) I-V relationship before subtraction; all I-V relationships were linear and almost identical in all solutions. (D3) I-V relationship of I_basal_ and I_evoked_ after Ba^2+^ subtraction. There were no clearly detectable currents, demonstrating the absence of any GIRK-likecurrents.

**Figure S2.**
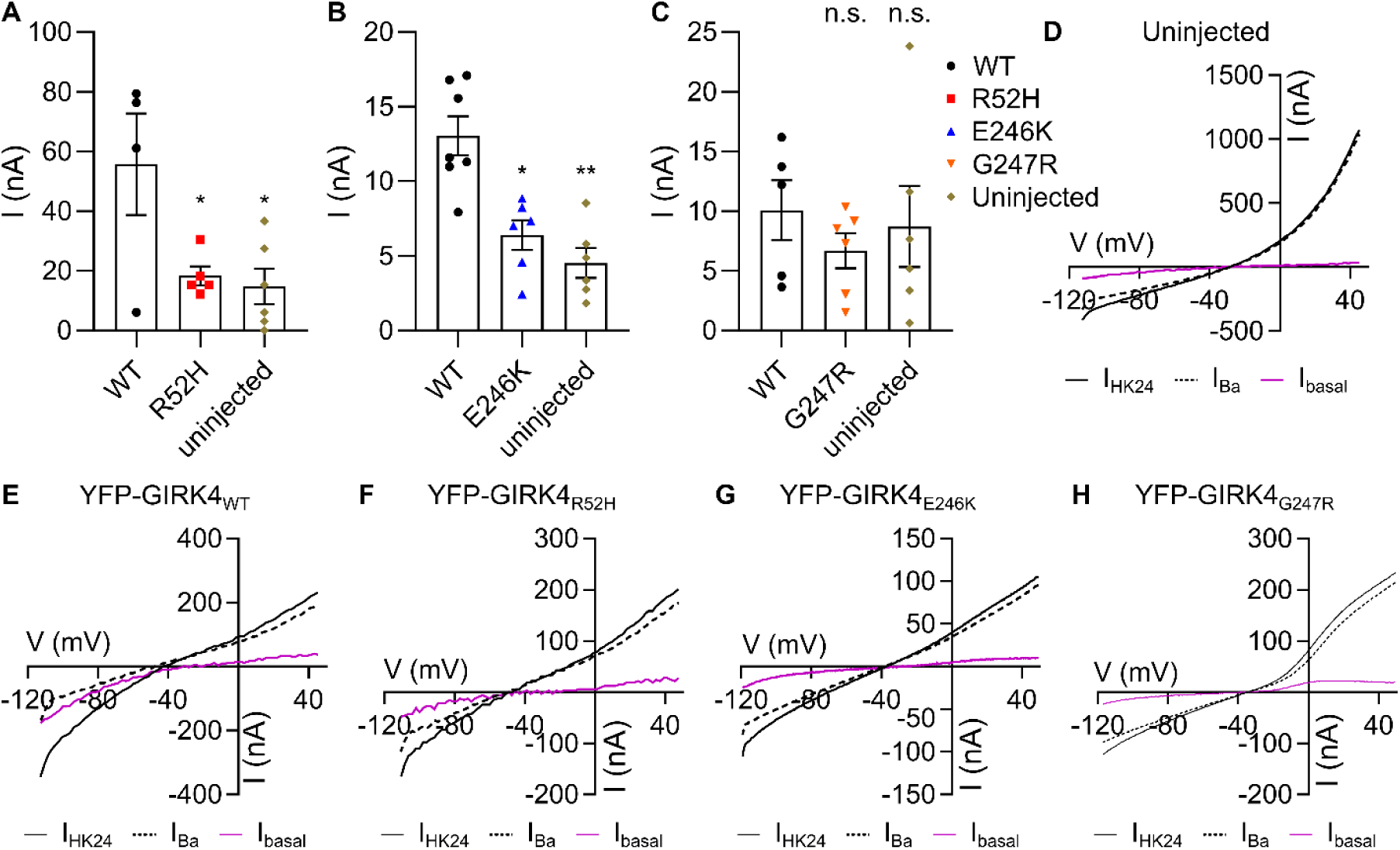
Homotetramers of YFP-GIRK4 channels are not active. (A-C) Currents in oocytes injected with RNAs of YFP-GIRK4 mutants, measured at −80 mV, were indistinguishable from currents recorded in uninjected oocytes. (D-H) Representative I-V relationships of uninjected oocytes (D) and homotetrameric YFP-GIRK4WT (E), YFP-GIRK4R52H (F), YFP-GIRK4E246K (G), YFP-GIRK4G247R (H).

**Figure S3.**
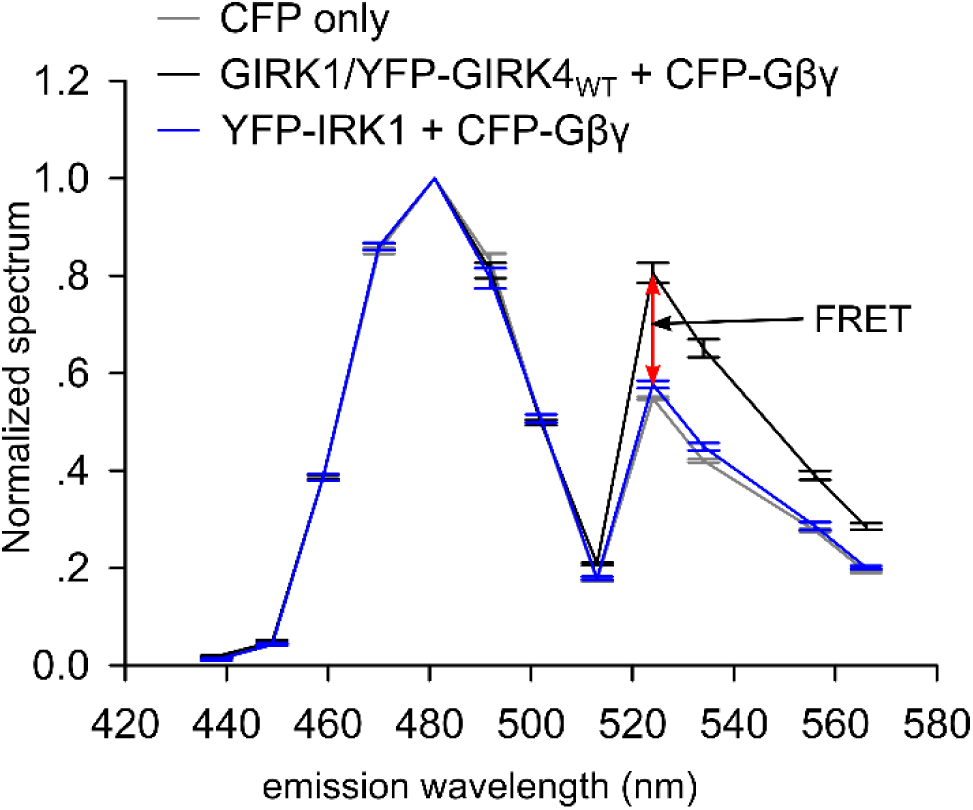
Emission spectra used to measure FRET. The graph shows emission spectra of GIRK1/YFP-GIRK4WT + CFP-Gβγ (black line), IRK1-YFP + CFP-Gβγ (light gray line) used as negative control, and CFP-Gβγ only (dark gray line), when the donor CFP was excited with the 405 nm laser. Values on X axis are midpoints of 11 nm-wide sampling windows in the spectral mode of Zeiss 501 Meta confocal microscope. GIRK1/YFP-GIRK4WT + CFP-Gβγ show large emission increase in YFP emission range (520 –575 nm) compared to the negative control and CFP only groups, demonstrating the energy transfer between the donor and acceptor molecules. Experimental points shown are mean±SEM from a representative experiment.

**Figure S4.**
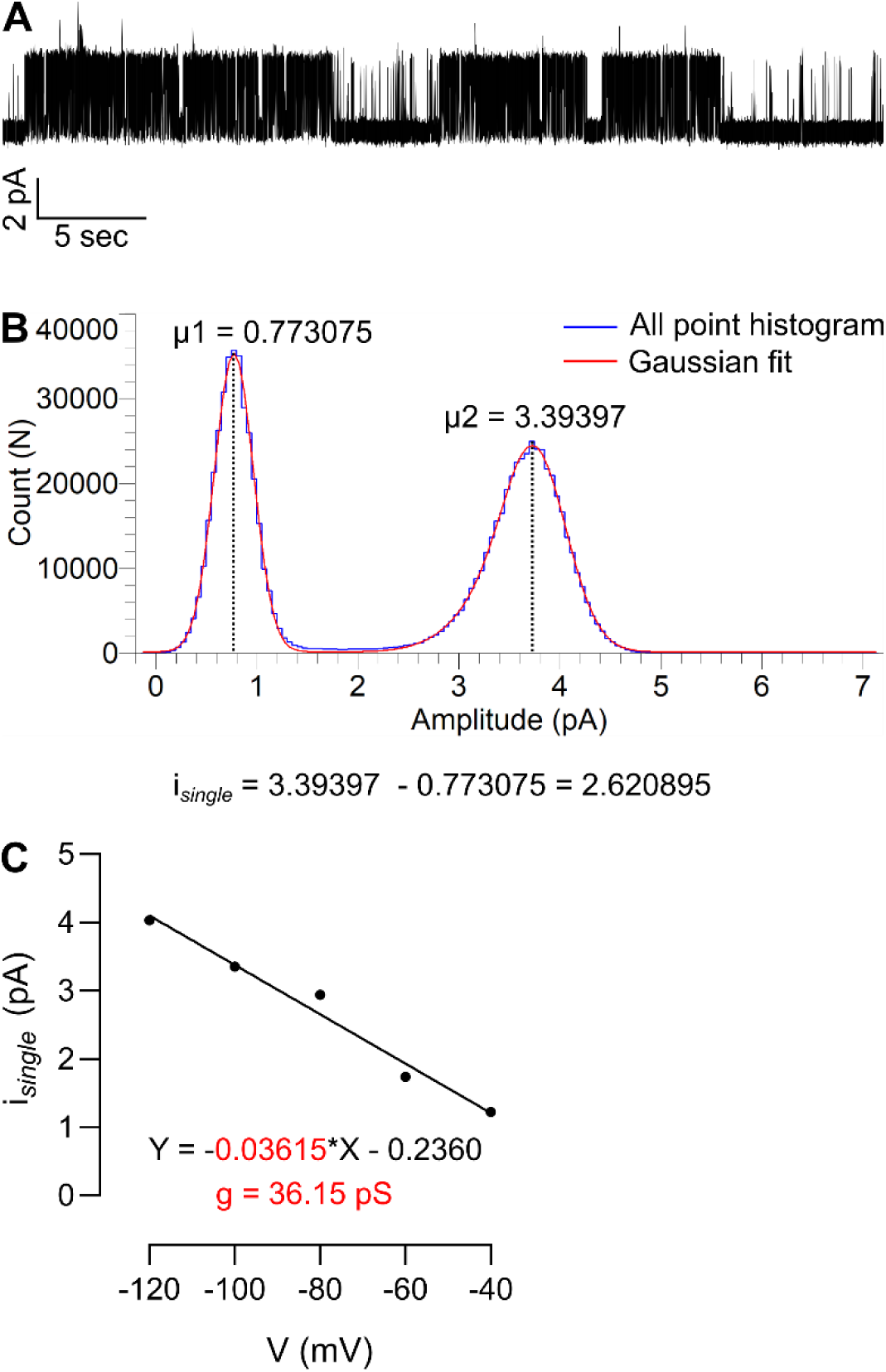
Estimation of i_*single*_ and single channel conductance from GIRK1/4 single channel recordings. (A) Example of a segment of the record that was used for the all point histogram in B. (B) All point histogram fitted with two Gaussians. The left peak is the background noise where the channel was closed, the right peak corresponds to the open state of the channel. Subtraction of the baseline (µ1) from the open state (µ2) gives i_*single*_. (C) Single channel conductance was derived from the slope of I-V relationship graph; i_*single*_ was calculated per each voltage in increments of 20 mV.

**Figure S5.**
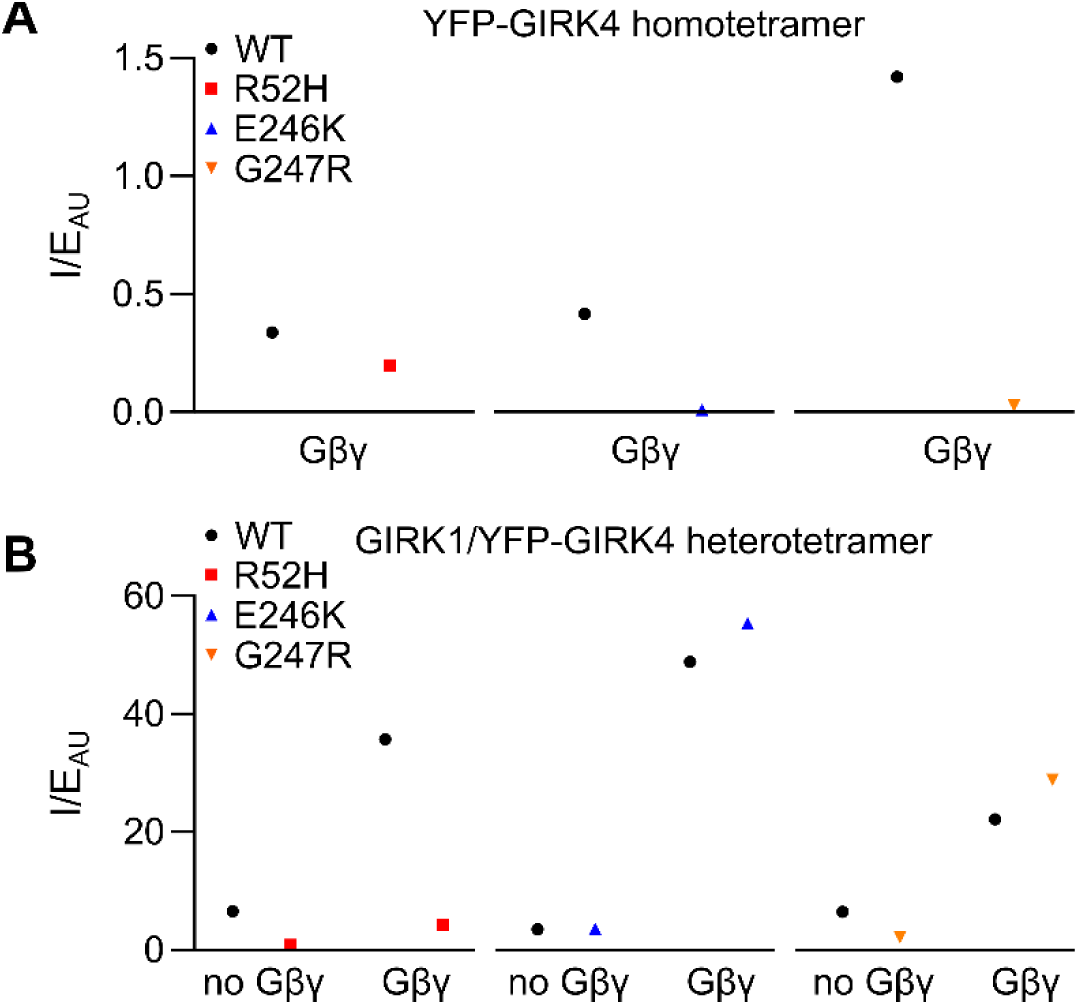
Analysis of whole-cell currents normalized to expression, for YFP-GIRK4 homotetramers and GIRK1/YFP-GIRK4 heterotetramers. (A) Normalization of current to expression of YFP-GIRK4 homotetramers indicates an impaired gating of the mutated channels. The normalization procedure included the division of mean value of current on mean value of channel expression in each experiment, giving one value for each experiment. Therefore, no statistical analysis could be performed. (B) Normalization of current to expression of GIRK1/YFP-GIRK4 heterotetramers indicates an impaired expression and gating of GIRK1/YFP-GIRK4R52H heterotetramers; on the other hand, E246K and G247R mutations appear to have the same open probability suggesting unimpaired gating.

## References

1. Logothetis DE, Kurachi Y, Galper J, Neer EJ & Clapham DE, The βγ subunits of GTP-binding proteins activate the muscarinic K^+^ channel in heart. Nature 325, 321–326 (1987).

2. Reuveny E, Slesinger PA, Inglese J, Morales JM, Iniguez-Lluhi JA, et al., Activation of the cloned muscarinic potassium channel by G protein βγ subunits. Nature 370, 143–146 (1994).

3. Krapivinsky G, Krapivinsky L, Wickman K & Clapham DE, Gβγ binds directly to the G protein-gated K^+^ channel, I_KACh_. J Biol Chem 270, 29059–29062 (1995).

4. Dascal N, Ion-channel regulation by G proteins. Trends Endocrinol Metab 12, 391–398 (2001).

5. Lu Z, Mechanism of rectification in inward-rectifier K^+^ channels. Annu Rev Physiol 66, 103–129 (2004).

6. Choi M, Scholl UI, Yue P, Bjorklund P, Zhao B, et al., K^+^ channel mutations in adrenal aldosterone-producing adenomas and hereditary hypertension. Science 331, 768–772 (2011).

7. Krapivinsky G, Gordon EA, Wickman K, Velimirovic B, Krapivinsky L, et al., The G-protein-gated atrial K^+^ channel I_KACh_ is a heteromultimer of two inwardly rectifying K^+^-channel proteins. Nature 374, 135–141 (1995).

8. Wickman K, Karschin C, Karschin A, Picciotto MR & Clapham DE, Brain localization and behavioral impact of the G-protein-gated K^+^ channel subunit GIRK4. J Neurosci 20, 5608–5615 (2000).

9. Wickman K, Nemec J, Gendler SJ & Clapham DE, Abnormal heart rate regulation in GIRK4 knockout mice. Neuron 20, 103–114 (1998).

10. Spat A & Hunyady L, Control of aldosterone secretion: a model for convergence in cellular signaling pathways. Physiol Rev 84, 489–539 (2004).

11. Hajnoczky G, Csordas G, Hunyady L, Kalapos MP, Balla T, et al., Angiotensin-II inhibits Na^+^/K^+^ pump in rat adrenal glomerulosa cells: possible contribution to stimulation of aldosterone production. Endocrinology 130, 1637–1644 (1992).

12. Korzyńska W, Jodkowska A, Goslawska K, Bogunia-Kubik K & Mazur G, Genetic aspects of primary hyperaldosteronism. Adv Clin Exp Med 27, 1149–1158 (2018).

13. Spat A, Glomerulosa cell—a unique sensor of extracellular K^+^ concentration. Mol Cell Endocrinol 217, 23–26 (2004).

14. Fernandes-Rosa FL, Boulkroun S & Zennaro MC, Somatic and inherited mutations in primary aldosteronism. J Mol Endocrinol 59, R47–R63 (2017).

15. Williams TA, Monticone S & Mulatero P, KCNJ5 mutations are the most frequent genetic alteration in primary aldosteronism. Hypertension 65, 507–509 (2015).

16. Monticone S, Tetti M, Burrello J, Buffolo F, De Giovanni R, et al., Familial hyperaldosteronism type III. J Hum Hypertens 31, 776–781 (2017).

17. Al-Salameh A, Cohen R & Desailloud R, Overview of the genetic determinants of primary aldosteronism. Appl Clin Genet 7, 67–79 (2014).

18. Hattangady NG, Karashima S, Yuan L, Ponce-Balbuena D, Jalife J, et al., Mutated *KCNJ5* activates the acute and chronic regulatory steps in aldosterone production. J Mol Endocrinol 57, 1–11 (2016).

19. Cheng CJ, Sung CC, Wu ST, Lin YC, Sytwu HK, et al., Novel *KCNJ5* mutations in sporadic aldosterone-producing adenoma reduce Kir3.4 membrane abundance. J Clin Endocrinol Metab 100, E155–163 (2015).

20. Murthy M, Xu S, Massimo G, Wolley M, Gordon RD, et al., Role for germline mutations and a rare coding single nucleotide polymorphism within the *KCNJ5* potassium channel in a large cohort of sporadic cases of primary aldosteronism. Hypertension 63, 783–789 (2014).

21. Hille B (2001) Ion Channels of Excitable Membranes (SINAUER ASSOCIATES, INC, Sunderland, Massachusetts U.S.A.).

22. Calloe K, Ravn LS, Schmitt N, Sui JL, Duno M, et al., Characterizations of a loss-of-function mutation in the Kir3.4 channel subunit. Biochem Biophys Res Commun 364, 889–895 (2007).

23. Whorton MR & MacKinnon R, X-ray structure of the mammalian GIRK2-βγ G-protein complex. Nature 498, 190–197 (2013).

24. Rubinstein M, Peleg S, Berlin S, Brass D, Keren-Raifman T, et al., Divergent regulation of GIRK1 and GIRK2 subunits of the neuronal G protein gated K^+^ channel by GαiGDP and Gβγ. J Physiol 587, 3473–3491 (2009).

25. Dascal N, The use of *Xenopus* oocytes for the study of ion channels. CRC Crit Rev Biochem 22, 317–387 (1987).

26. Dascal N, Landau EM & Lass Y, *Xenopus* oocyte resting potential, muscarinic responses and the role of calcium and guanosine 3’,5’-cyclic monophosphate. J Physiol 352, 551–574 (1984).

27. Monticone S, Hattangady NG, Penton D, Isales CM, Edwards MA, et al., A novel Y152C *KCNJ5* mutation responsible for familial hyperaldosteronism type III. J Clin Endocrinol Metab 98, E1861–1865 (2013).

28. Zheng J, Varnum MD & Zagotta WN, Disruption of an intersubunit interaction underlies Ca2+-calmodulin modulation of cyclic nucleotide-gated channels. J Neurosci 23, 8167–8175 (2003).

29. Berlin S, Tsemakhovich VA, Castel R, Ivanina T, Dessauer CW, et al., Two distinct aspects of coupling between Gαi protein and G protein-activated K^+^ channel (GIRK) revealed by fluorescently labeled Gαi3 protein subunits. J Biol Chem 286, 33223–33235 (2011).

30. Bykova EA, Zhang XD, Chen TY & Zheng J, Large movement in the C terminus of CLC-0 chloride channel during slow gating. Nat Struct Mol Biol 13, 1115–1119 (2006).

31. Chen AX, Nishimoto K, Nanba K & Rainey WE, Potassium channels related to primary aldosteronism: Expression similarities and differences between human and rat adrenals. Mol Cell Endocrinol 417, 141–148 (2015).

32. Aragao-Santiago L, Gomez-Sanchez CE, Mulatero P, Spyroglou A, Reincke M, et al., Mouse Models of Primary Aldosteronism: From Physiology to Pathophysiology. Endocrinology 158, 4129–4138 (2017).

33. Kozek KA, Du Y, Sharma S, Prael FJ, 3rd, Spitznagel BD, et al., Discovery and characterization of VU0529331, a synthetic small-molecule activator of homomeric G protein-gated, inwardly rectifying, potassium (GIRK) channels. ACS Chem Neurosci 10, 358–370 (2019).

34. Murthy M, Azizan EA, Brown MJ & O’Shaughnessy KM, Characterization of a novel somatic *KCNJ5* mutation delI157 in an aldosterone-producing adenoma. J Hypertens 30, 1827–1833 (2012).

35. Kuppusamy M, Caroccia B, Stindl J, Bandulik S, Lenzini L, et al., A novel *KCNJ5*-insT149 somatic mutation close to, but outside, the selectivity filter causes resistant hypertension by loss of selectivity for potassium. J Clin Endocrinol Metab 99, E1765–1773 (2014).

36. Hardege I, Xu S, Gordon RD, Thompson AJ, Figg N, et al., Novel insertion mutation in *KCNJ5* channel produces constitutive aldosterone release from H295R cells. Mol Endocrinol 29, 1522–1530 (2015).

37. Kokunai Y, Nakata T, Furuta M, Sakata S, Kimura H, et al., A Kir3.4 mutation causes Andersen-Tawil syndrome by an inhibitory effect on Kir2.1. Neurology 82, 1058–1064 (2014).

38. Kuß J, Stallmeyer B, Goldstein M, Rinne S, Pees C, et al., Familial sinus node disease caused by a gain of GIRK (G-protein activated inwardly rectifying K^+^ channel) channel function. Circ Genom Precis Med 12, e002238 (2019).

39. Charmandari E, Sertedaki A, Kino T, Merakou C, Hoffman DA, et al., A novel point mutation in the *KCNJ5* gene causing primary hyperaldosteronism and early-onset autosomal dominant hypertension. J Clin Endocrinol Metab 97, E1532–1539 (2012).

40. Bukiya AN & Rosenhouse-Dantsker A, Synergistic activation of G protein-gated inwardly rectifying potassium channels by cholesterol and PI(4,5)P_2_. Biochim Biophys Acta Biomembr 1859, 1233–1241 (2017).

41. Vivaudou M, Chan KW, Sui JL, Jan LY, Reuveny E, et al., Probing the G-protein regulation of GIRK1 and GIRK4, the two subunits of the K_ACh_ channel, using functional homomeric mutants. J Biol Chem 272, 31553–31560 (1997).

42. Doyle DA, Morais Cabral J, Pfuetzner RA, Kuo A, Gulbis JM, et al., The structure of the potassium channel: molecular basis of K^+^ conduction and selectivity. Science 280, 69–77 (1998).

43. Lu Q & Miller C, Silver as a probe of pore-forming residues in a potassium channel. Science 268, 304–307 (1995).

44. Inanobe A, Matsuura T, Nakagawa A & Kurachi Y, Structural diversity in the cytoplasmic region of G protein-gated inward rectifier K^+^ channels. Channels (Austin) 1, 39–45 (2007).

45. Nishida M & MacKinnon R, Structural basis of inward rectification: cytoplasmic pore of the G protein-gated inward rectifier GIRK1 at 1.8 A resolution. Cell 111, 957–965 (2002).

46. Pegan S, Arrabit C, Zhou W, Kwiatkowski W, Collins A, et al., Cytoplasmic domain structures of Kir2.1 and Kir3.1 show sites for modulating gating and rectification. Nat Neurosci 8, 279–287 (2005).

47. Kubo Y & Murata Y, Control of rectification and permeation by two distinct sites after the second transmembrane region in Kir2.1 K^+^ channel. J Physiol 531, 645–660 (2001).

48. Hommers LG, Lohse MJ & Bunemann M, Regulation of the inward rectifying properties of G-protein-activated inwardly rectifying K^+^ (GIRK) channels by Gβγ Subunits. J Biol Chem 278, 1037–1043 (2003).

49. Hedin KE, Lim NF & Clapham DE, Cloning of a *Xenopus laevis* inwardly rectifying K^+^ channel subunit that permits GIRK1 expression of I_KACh_ currents in oocytes. Neuron 16, 423–429 (1996).

50. Dascal N & Lotan I (1992) in. Protocols in Molecular Neurobiology, eds. Longstaff, & Revest, P. (Springer New York, Totowa, NJ), pp. 205–225.

51. Rubinstein M, Peleg S, Berlin S, Brass D & Dascal N, Gαi3 primes the G protein-activated K^+^ channels for activation by coexpressed Gβγ in intact *Xenopus* oocytes. J Physiol 581, 17–32 (2007).

52. Parmar J, Key RE & Rainey WE, Development of an adrenocorticotropin-responsive human adrenocortical carcinoma cell line. J Clin Endocrinol Metab 93, 4542–4546 (2008).

53. Raifman TK, Kumar P, Haase H, Klussmann E, Dascal N, et al., Protein kinase C enhances plasma membrane expression of cardiac L-type calcium channel, CaV1.2. Channels (Austin) 11, 604–615 (2017).

54. Yakubovich D, Berlin S, Kahanovitch U, Rubinstein M, Farhy-Tselnicker I, et al., A quantitative model of the GIRK1/2 channel reveals that its basal and evoked activities are controlled by unequal stoichiometry of Gα and Gβγ. PLoS Comput Biol 11, e1004598 (2015).

55. Yakubovich D, Rishal I, Dessauer CW & Dascal N, Amplitude histogram-based method of analysis of patch clamp recordings that involve extreme changes in channel activity levels. J Mol Neurosci 37, 201–211 (2009).

56. Sakmann B & Neher E (1995) Single-Channel Recording.

57. Kanevsky N & Dascal N, Regulation of maximal open probability is a separable function of Ca_vβ_ Subunit in L-type Ca^2+^ channel, dependent on NH_2_ terminus of α_1C_ (Ca_v_1.2α). J Gen Physiol 128, 15–36 (2006).

